# LevSeq: Rapid Generation of Sequence-Function Data for Directed Evolution and Machine Learning

**DOI:** 10.1101/2024.09.04.611255

**Authors:** Yueming Long, Ariane Mora, Emre Gürsoy, Kadina E. Johnston, Francesca Zhoufan-Li, Frances H. Arnold

## Abstract

Sequence-function data provides valuable information about the protein functional landscape, but is rarely obtained during directed evolution campaigns. Here, we present Long-read every variant Sequencing (LevSeq), a pipeline that combines a dual barcoding strategy with nanopore sequencing to rapidly generate sequence-function data for entire protein-coding genes. LevSeq integrates into existing protein engineering workflows and comes with open-source software for data analysis and visualization. The pipeline facilitates data-driven protein engineering by consolidating sequence-function data to inform directed evolution and provide the requisite data for machine learning-guided protein engineering (MLPE). LevSeq enables quality control of mutagenesis libraries prior to screening, which reduces time and resource costs. Simulation studies demonstrate LevSeq’s ability to accurately detect variants under various experimental conditions. Finally, we show LevSeq’s utility in engineering protoglobins for new-to-nature chemistry. Widespread adoption of LevSeq and sharing of the data will enhance our understanding of protein sequence-function landscapes and empower data-driven directed evolution.

## Introduction

Directed evolution (DE) has been key to the discovery and engineering of biocatalysts for new-to-nature chemistry^1^, development of sustainable bioprocesses for pharmaceutical synthesis^2,3^, and for engineering proteins for environmental sensing^4^ and bioremediation^5^, among many other applications. The power of directed evolution resides in the rapid evaluation of mutated proteins to traverse the fitness landscape toward those exhibiting improved fitness^6,7^. A typical directed evolution campaign involves the generation and screening of thousands of variants – a significant number but still only a tiny fraction of the possible sequence space^8^. To streamline directed evolution, machine learning (ML) can be employed^9–11^ to guide sequence-function exploration to variants with high fitness^12–15^.

Traditional directed evolution (DE) approaches have generated datasets rich in activity labels but often lacking sequence information, as they focus on optimizing activity without sequencing all variants^3^. Existing sequence-function datasets for protein evolution studies are primarily comprised of deep mutational scanning data covering all single mutations or combinatorial libraries targeting specific sites^16–18^. While valuable, these approaches are costly and capture only a fraction of the sequence diversity most useful for protein evolution^19^. To advance machine learning in protein engineering, we need a method for collecting, analyzing, and pairing sequence-function data from diverse mutagenesis approaches^20,21^. This method would work for random mutagenesis across whole genes, combinatorial libraries at sites distant in the primary sequence, and other targeted mutagenesis approaches.

Challenges that must be overcome to realize this vision include the high cost of sequencing entire genes^22^ and the lack of a standardized format to create and distribute the data. The Arnold lab developed the every variant sequencing (evSeq) method using Illumina short-read sequencing to capture the sequences of variants arrayed in 96-well plates^23^. Due to the short sequencing lengths (∼250 base pairs), however, evSeq is not ideal for collecting full-gene-length gene sequences. In contrast, real-time sequencing technologies like nanopore sequencing can capture millions of long reads at a low cost^24^, but nanopore sequencing is characterized by a high error rate^25–27^. Previously published high-throughput nanopore sequencing methods, parSEQ^28^ and SequenceGenie^27^, have overcome this limitation by performing statistical analyses on consensus reads to detect true variants. UMIC-seq takes a different approach and clusters sequences rather than identifying individual variants, as the objective is to map evolutionary lineages^29^. Each of these methods uses a similar DNA-barcoding approach to demultiplex reads, which we have now coupled with the evSeq pipeline to enable collection of sequence-function data for directed evolution studies.

This work describes Long-read every variant Sequencing (LevSeq), which extends the previous evSeq method by utilizing the barcode strategy described in Currin et al.^27^ for nanopore sequencing, enabling the evSeq pipeline to be utilized on full-length genes. LevSeq includes the following steps: 1) a colony polymerase chain reaction (PCR) to generate barcoded gene amplicons, 2) Oxford Nanopore sample preparation and sequencing to generate sequencing information, 3) demultiplexing and variant identification, 4) sequence-function data coupling accompanied by visualization and analyses, and 5) generation of data outputs that are amenable to downstream ML and compatible with existing databases. This method is rapid and robust under different mutagenesis conditions and enables researchers with no prior experience working with next-generation sequencing (NGS) data to perform analyses of mutagenesis libraries.

Importantly, LevSeq offers several advantages: a) the software is open source, easy to set up, and designed for directed evolution experiments; b) it requires as few as ten reads to detect a variant in a well; c) results are available before the resource-intensive screening phase, enabling selection of specific variants for testing; d) fitness data are linked with sequence information to inform subsequent engineering steps. We demonstrate LevSeq in two protein engineering projects. First, we sequenced ∼1,000 variants of an error-prone polymerase chain reaction (epPCR) random mutagenesis library and identified the top variants in a typical epPCR workflow by coupling sequence and function data. In the second demonstration, we applied LevSeq to variants sampled from a five-site combinatorial library, which yielded data for downstream ML packages to predict variants with increased activity^30^. We show that LevSeq facilitates machine learning-guided protein engineering (MLPE) by collecting a small subset of sequence-function data from the studied system to be used as training or input data.

## Materials and Methods

### Design of backbone-specific barcoded primers

Universal binding sites of the pET-22b(+) cloning vector are first identified, and two sites upstream and downstream of the cloning site were chosen for primer design. All variants cloned using the pET-22b(+) vector can be sequenced using the primers designed for this research. Alternatively, primers can be designed for different cloning vectors, as long as the barcodes are attached to the upstream of the upstream primer and downstream of the downstream primer. (Supplementary Information Oligonucleotide Design).

### Colony PCR for generating barcoded amplicons

PCR protocols are optimized for robust amplification of the full-length gene. Best performance is obtained using Taq polymerase and a touchdown PCR program. The PCR set up for each well includes 1 µL of overnight culture, 2 µL of 1 µM each barcoded primer mix, and 7 µL of PCR master mix. Using either 96-well or 384-well PCR thermocyclers, an initial 300 s denaturing step is performed followed by the touchdown PCR program detailed in the supplementary information. Depending on the length of the gene, one minute elongation time per 1 kb is recommended for optimal amplification. Amplicons were then pooled (Supplementary Information), analyzed, and purified with gel electrophoresis using the Zymoclean Gel DNA Recovery Kit (Zymo Research D4002).

### Sample preparation and sequencing

Purified amplicon samples were normalized and combined into one sample and prepared for sequencing using the Oxford Nanopore ligation sequencing kit (LSK-114). For the MinION and Flongle run setup, 0.02 Gb of basecalled bases per 96 variants and super accurate basecalling model are recommended as sequencing parameters. One Flongle flow cell is recommended for sequencing up to 1600 variants, whereas the MinION flow cell can be washed and reused using the Oxford Nanopore wash kit until the number of pores decreases below 100. We recommend skipping the use of storage buffer as significant pore loss was observed after applying storage buffer to the flow cell (∼300 pores lost).

### Variant calling

The computational delineation of reads from a pooled sample is slow and thus we wrote a bespoke pairwise local alignment using the Smith-Waterman algorithm in C++ to efficiently detect barcodes at the 5’ and 3’ ends of each nanopore read. The 3’ end barcodes are aligned to the last 100 base pairs of each read, and the highest matching score above threshold 80 is used to assign each read to the 96-well plate of origin. Next, 5’ barcodes are aligned to the first 100 base pairs of each read, and the highest matching score is used to assign each read to a specific well within the assigned plate. The reads for each well are aligned using minimap2, version 2.1. The parameters for minimap2 are the standard long read parameters: “-ax map-ont”, a -B mismatch score of 2, a match score of 4, and a gap opening penalty of 10. These were chosen to deprioritize frame shift mutations, because they occur less frequently. If multiple reads with the same read ID are mapped to the same well, the read with the highest quality is retained. During variant calling several quality control files are produced: a multiple sequence alignment and a csv file for each well in each plate, which contains the p-values, p-adjusted values, and counts for each position in the sequence. The sequencing error for a nanopore device is comparatively high at approximately 10% and dependent on myriad factors such as the flow cell, age of the cell, run conditions, etc. As such, for each well we calculate the error rate as the mean rate of non-reference nucleic acids per position. The probability that a mutation observed across a set of reads is a true mutation can be calculated using the binomial test. Namely, the null hypothesis, *π*_0_, is that the observed sequence variation is due to the inherent sequencing error of the nanopore device, and the alternative hypothesis is that the observed nucleic acids are due to a mutation induced by SSM or epPCR. The number of trials, *n*, is the number of reads for a given well, the number of successes, *k*, is the number of a given nucleic acid or deletion, that is different to the reference sequence. Significance of the observed data is calculated using a one-sided test, testing for greater than expected error *π* > *π*_0_, and calculating this for each A, T, G, C and deletion for each position for each well. The expected error rate used can be defined by the user; we use a default of 10%, or the mean error rate for the well. Multiple testing is corrected for by using the Benjamini-Hochberg test, with a false discovery rate of 0.05. For each well, for a given sequence, the number of tests that are corrected for is equivalent to the length of the sequence. Additionally, corrections for multiple testing are made across the wells that meet a mutation frequency threshold. If a well has a mutation frequency above a user-defined threshold, the well is checked for mutations and mixed wells. A well is classified as “mixed” if a position has more than one significant mutation by the FDR adjusted binomial test, using a threshold of p < 0.05 by default. Finally, post-variant calling we match the nucleotide variants to the amino acid changes.

### Simulation study

For the simulation, the protoglobin used in case 1 and case 2 was chosen. This protein is 204 amino acids long. For epPCR, errors are introduced at the DNA level, and as such an error rate of 2% corresponds to approximately 12 nucleotide mutations. To test the effect of sequencing error, sequencing error was varied from 0 to 100% across the sequence by incrementing at 5% intervals with a constant read depth of 10 reads. For read depth, the number of reads varied from one read to 50 in increments of 1, with the sequencing error held constant at the reported nanopore sequencing error of 10%. To test the effect of sequence length, the sequence was trimmed to lengths between five and 200 at step sizes of 20.

### Software stack

After the initial base-calling, which is a default option when using the MinION protocol, LevSeq is operating-system independent and runs entirely using docker or a web app. We modularized the application to comprise two components. The first is a command line and app that is used to de-multiplex and call variants. This is hosted locally, to reduce the need for large data transfer and processing. We opted to deploy this as an in-house tool as this component is stable and unlikely to change in future software updates. The code and docker image are available on GitHub (https://github.com/fhalab/LevSeq). It includes a multiple sequence alignment that shows the pileup from the bam files for quality control visualization. The output is an interactive HTML file that provides a per-plate view of the mutation, sequence count, and alignment probability. The second component is a web application where users upload screening data with coupled function. To calculate the combined “fitness”, we normalize each user-provided feature and then compute the median across the normalized features. While median is the recommended summary statistic, as it is less susceptible to outliers, users can switch to calculating the mean across features.

### Error-prone PCR random mutagenesis library generation for ParLQ

Error-prone mutagenesis libraries were prepared using a standard error-prone PCR protocol. We designed primers using the template given in Supporting Information, Table S13. Different concentrations (100 mM, 200 mM, 300 mM, 400 mM) of MnCl_2_ were added to each PCR. Once PCRs finished, 1 µL of DpnI (NEB R0176S) was added to each of the reactions followed by incubation at 37 ^°^C for 1 h to digest any residual template plasmid. DNA fragments with the desired size were excised from an agarose electrophoresis gel and then purified using the Zymoclean Gel DNA Recovery Kit (Zymo Research D4002).

The expression plasmids containing an ampicillin resistance gene were constructed following standard Gibson assembly method. After 1 h of incubation at 50 ^°^C, the reaction mixtures were used to transform T7 Express competent BL21 *E. coli* cells (NEB C2566H). Transformed cells were spread onto solid agar selection medium consisting of Luria broth (RPI L24040-5000.0) supplemented with 0.1 mg/mL ampicillin (LBamp) and incubated at 30 ^°^C until visible individual colonies are formed. To grow the error-prone libraries, 600 µL of LBamp were added into each well of 96-well deep-well plates (2-mL well volume). Individual colonies from the agar plates were then transferred into the wells with each well containing a single colony. The plates containing these overnight cultures were shaken at 220 rpm, 37 ^°^C, and 80% humidity for 16 hours in an Infors Multitron HT shaking incubator. After overnight growth, 100 µL of overnight cultures were added to 100 µL of 50% glycerol solution to make glycerol stock plates, these plates can be used to store variants for future analysis.

### Sequencing of ParLQ epPCR libraries

With the fresh overnight culture, sequencing libraries were prepared following the protocol described in Supporting Information, LevSeq Library Preparation and Sequencing; the LevSeq software was run using all default parameters. Barcode-linked primer plates used are in Supporting information, Tables S5–S12; the barcode plates were paired to libraries as given in Supporting Information, Table S14.

### Measuring *cis* and *trans* cyclopropane formation from 4-methoxystyrene

For expression of the variant libraries, 50 µL of the saturated overnight cultures were used to inoculate 900 µl of Terrific Broth with 0.1 mg/mL ampicillin (TBamp) in 96-well plates. These cultures were then grown at 37 ^°^C, 220 rpm, and 80% humidity for 2.5 h in an Infors Multitron HT shaking incubator, after which they were placed on ice for 20 minutes. Following this, 25 µL of a 20 mM solution of isopropyl-*β*-d-thiogalactoside (IPTG; GoldBio # I2481C100) and 25 µL of a 40 mM solution of 5-aminolevulinic acid (ALA;thermo scientific # 103920050) in TBamp were added to each well to induce protein expression at a final concentration of 0.5 mM IPTG and 1 mM ALA. Expression proceeded in the same Infors shaker at 22 ^°^C and 220 rpm for 18 h. Cells were harvested through centrifugation at 4,000g for 5 minutes, the supernatant was removed, and the pellets were resuspended in 380 µL of M9-N. In an anaerobic Coy chamber, 20 µL of 200 mM 4-methoxystyrene and 300 mM ethyldiazoacetate in acetonitrile were added into each well of the resuspended pellets. The reaction plates were sealed using sticky aluminum foil and shaken at room temperature at 800 rpm on an IKA MTS 4 shaker for 20 h. Following the reaction, 800 *µ*L of cyclohexane were added to each well, and the reactions were shaken and centrifuged at 5,000g for 10 minutes. The organic layer was transferred into GC screw vials (Agilent 5182-0715) and analyzed using GCMS (Agilent 7820A(G4350A)).

## Results and Discussion

### A standardized workflow to sequence thousands of full-length variant genes

We use a dual backbone-specific barcoded primer system to streamline the sequencing process and maximize resource efficiency^31^. The primer sequences are designed for pET-22b(+) backbones and can be redesigned for other cloning vectors following standard design techniques (Supplementary Information Oligonucleotide Design). Compared to the original evSeq approach, LevSeq is not constrained by sequence length, can sequence any gene of interest in the cloning backbone, and has a short turnaround time (3–12 hours)^23^. The protocol commences with a one-step colony PCR that produces a full-length protein-coding DNA amplicon with unique barcode pairs at both ends (Figure 1A). Using 96 unique forward barcodes for each well of a 96-well plate and 96 unique reverse barcodes for as many as 96 unique plates, it is theoretically possible to demultiplex and sequence 9,216 variants^29–32^.

**Figure 1:**
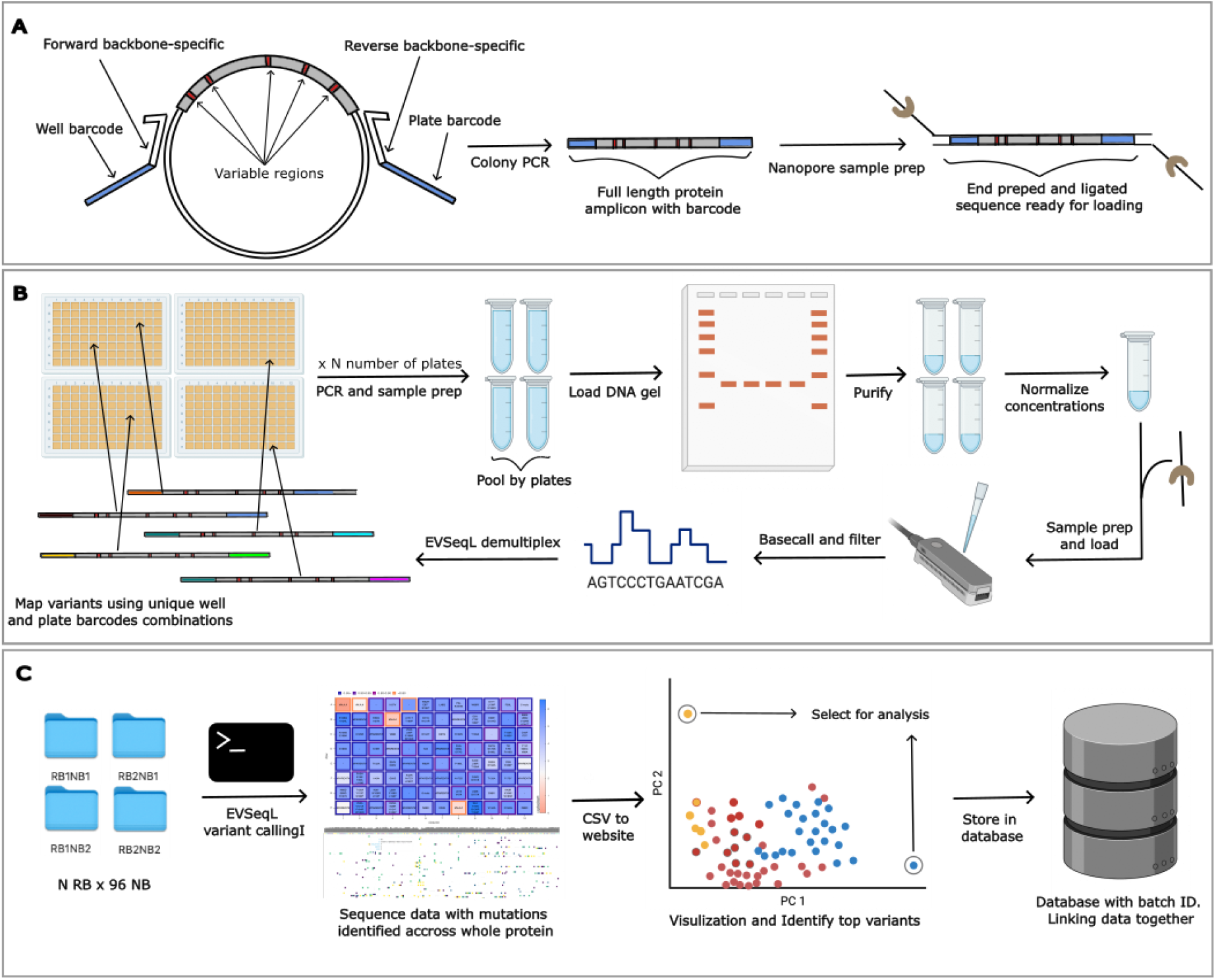
Overview of LevSeq library preparation, variant sequencing, and data visualization. A. The first step of LevSeq involves a one-step PCR using backbone-specific 5’ and 3’ end barcoded primers to amplify the full-length targeted gene. B. All PCR products from one 96-well plate are pooled for gel purification. The purified DNA samples from each plate are normalized by molarity and combined for nanopore sample preparation using the ligation sequencing kit. The sequencing run is performed in-house on a MinION sequencer, and the raw voltage signals are base-call-converted into nucleotides with the resulting fastq reads filtered by quality. C. Sequence function data pairing, visualization, and storage in format compatible with database.

A typical round of directed evolution with LevSeq begins with isolating colonies into a 96-well plate for overnight culture, followed by protein expression and screening. The LevSeq protocol is executed during the protein expression time, after overnight cultures of individually arrayed colonies in 96-well plates are grown to saturation.

A small amount of overnight culture is combined with PCR master mix and barcoded primers to generate barcoded gene amplicons from each well (Supplementary Protocols). After colony PCR, DNA samples are pooled and normalized (Figure 1B). Samples are prepared for sequencing using the ligation kit provided by Oxford Nanopore before being loaded onto the MinION or Flongle flow cell. The real-time sequencing data, stored as raw and base-called files, are readily processed by the open-source software for comprehensive data analysis, visualization, and storage (Figure 1C). Our software also provides a template for LC-MS instruments based on the sequencing results to screen only true variants, reducing the screening load and automatically coupling the sequence function data (Figure 2).

**Figure 2:**
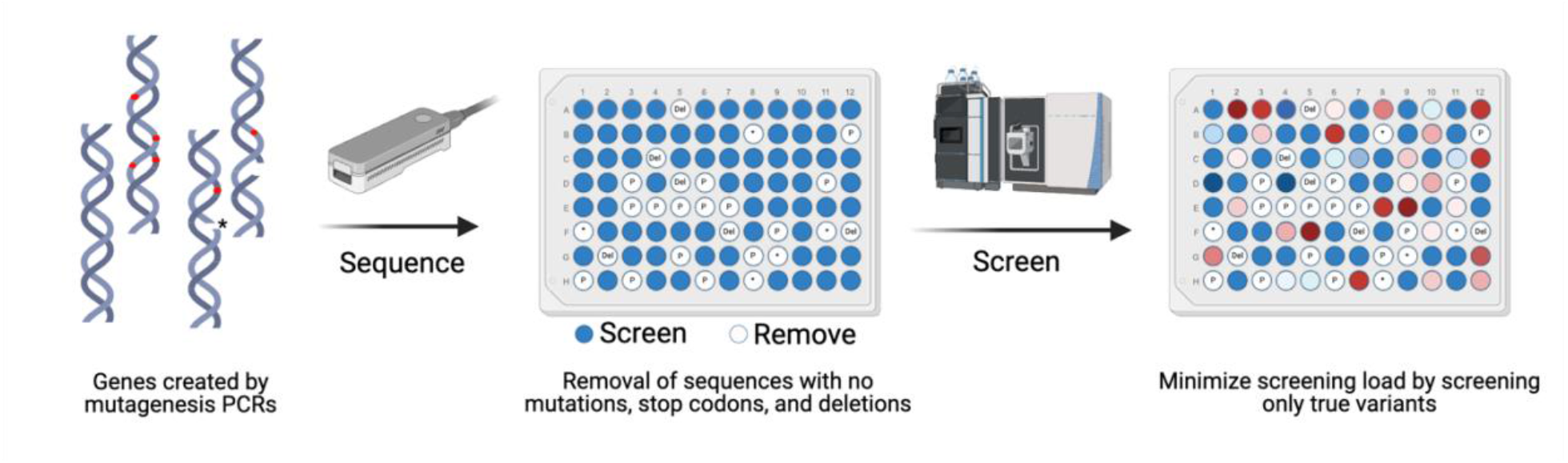
LevSeq reduces screening burden by enabling removal of sequences with no mutations, stop codons, and deletions.

### LevSeq has an accurate variant caller that is robust to experimental designs

We developed a software suite for LevSeq that consists of two components to process and analyze every variant. The first is an operating system-independent docker image that performs efficient barcode de-multiplexing and runs on the sequencing computer^36^. The de-multiplexed plate and well data are parsed by a Python package to identify statistically significant variants and notify users of any poor-quality mappings or mixed wells. If a particular barcode is undesirable, users can edit the barcode sequences file provided in the software suite to incorporate any customized barcodes; no information beyond the barcode sequence is needed for the demultiplexing step. We validated the variant calling software by performing over 1,000 simulations to test the effect of experimental and sequencing conditions on the variant calling accuracy. Experimental variation, defined here as protein sequence length and ePCR error rate showed no effect on the efficacy of variant calling (Supplementary Figure 1E–F). Sequencing variation, such as nanopore error rate, does not affect the ability to accurately call variants if more than 10 reads are assigned to a well, which is within the typical flow cell operating range (error < 20%), (Supplementary Figure 1A–B). We showed that variants are accurately detected (>99%) with 10 reads^37^ to generate a consensus sequence^38,39^ up to a sequencing error rate of 20%, which exceeds the expected error of nanopore devices^40^ (Figure 3A). Previous methods have opted for alternative approaches to overcome the high error rate: SequenceGenie leverages strand bias and Bayesian statistics to identify true variants^27^, while parSEQ uses a conservative consensus threshold of 0.9^28^.

**Figure 3:**
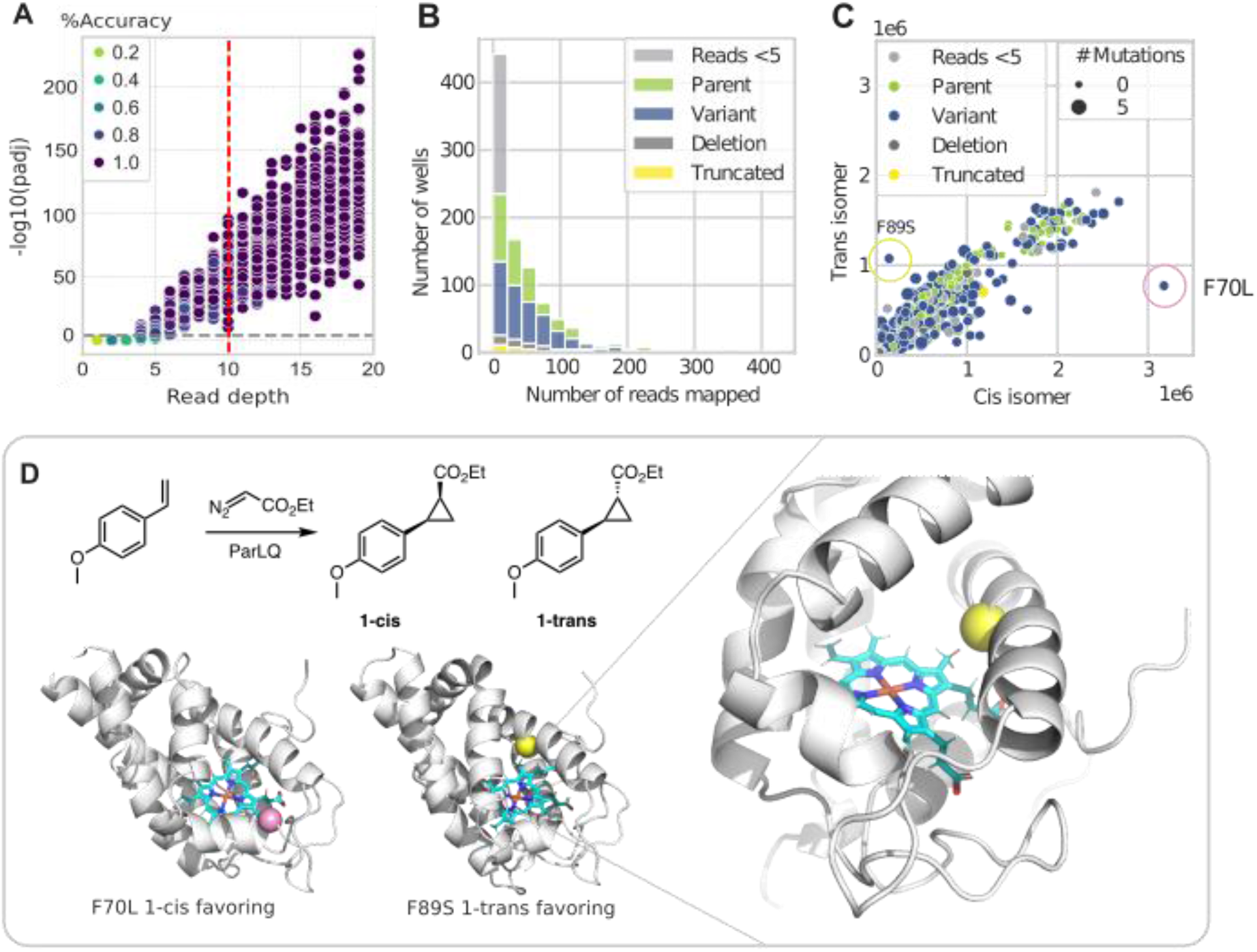
A. Accuracy of detecting variants using simulation studies on ParPgb LQ, varying read depth from 1 to 20 using an epPCR mutation rate of 2% and a nanopore sequencing error rate of 10%. B. Beneficial epPCR mutations favoring the *cis* and *trans* products are shown on the ParPgb structure. C. Clustered reads across all ten plates from the epPCR experiment. D. Enzyme-catalyzed reaction screened in the epPCR experiments is cyclopropanation of 4-methoxystyrene, leading to *cis* and *trans* cyclopropane products. The positions mutated for selected variants are highlighted on an AlphaFold3 structure of ParPgb.

The second component of the software suite is a website that processes LevSeq files for future reference and downstream ML. Users upload variant information from LevSeq along with associated fitness data. Top variants are returned to the user along with a visualization of the coupled sequence function data. This process enables the collection and consolidation of standardized data along with the associated screening conditions (e.g. chemical reaction, stability, etc.). The deployment of this software is a primary differentiator between LevSeq and other nanopore variant calling methods, parSEQ and SequenceGenie. While SequenceGenie is also suitable for an individual lab, it requires users to build the docker image and has limited documentation, making it a challenge for the standard bench scientist to implement^27^. ParSEQ is an extensive software suite and utilizes comprehensive cloud computing, making it ideal for larger scale operations. However, it requires knowledge of cloud infrastructure^28^, an uncommon skill set in a typical protein engineering lab. LevSeq followed the evSeq approach and was designed to be easily deployable with minimal installation and a single command to run analyses and output data in an interoperable format. We envision these datasets will be compatible with existing databases^41–44^ and will become a useful resource for protein engineers who seek to create data-driven models of protein sequence-function landscapes.

### Use case 1: Analyze random mutagenesis libraries and inform next steps in DE

To demonstrate the utility of LevSeq in a random mutagenesis experiment, we constructed and screened ten 96-well plates from an error-prone PCR library of *Pyrobaculum arsenaticum* protoglobin (*Par*Pgb) LQ variants^45^. *Par*Pgb is isolated from thermostable archaea and expresses well in *Escherichia coli*. Over the past decade, our laboratory has shown that protoglobins exhibit remarkable tolerance to mutations that alter their catalytic capabilities^46–49^.

Three of the ten plates exhibited unusually low sequencing coverage, contributing to a large number of samples with insufficient coverage (Supplementary Figure 2). (The insufficient coverage in this case resulted from improper PCR amplification. This can be mitigated by adjusting the PCR setup method (Supplementary Protocols) and was not observed in subsequent sequencing experiments (Supplementary Figure 3). The final dataset for *Par*Pgb included 211 sequences with zero amino acid mutations and 539 sequences with up to five amino acid mutations from the parent. Single amino acid mutations were most prevalent, occurring in 210 out of 539 sequences (Figure 3B and 3C). The mutation distribution aligned with the expected outcome of the mutagenesis method. Following sequencing, we utilized the LevSeq toolkit to generate sequence-function data.

All ten plates were screened for activity and specificity for catalyzing the formation of the *cis* and *trans* cyclopropanation products of 4-methoxystyrene (Figure 3A). The software automatically returns the top-performing variants for each recorded fitness value, which in this case were top variants for both the *cis* and *trans* cyclopropanation products (see Methods for details on selection criteria). We found a single mutation conferred the highest activity for each desired stereoisomeric outcome: F70L for *cis* preference and F89L for *trans* (Figure 3D). Sequencing every variant also revealed sites with epistatic interactions. For example, the single mutation D72G improved the formation of both *cis* and *trans* products 1.5-fold, and the F89L mutation improved *trans* product formation nearly 3-fold. However, combining D72G and F89L resulted in activity similar to the parent, indicating higher-order interactions^50,51^ between sites 72 and 89, which could be further investigated with a double-site saturation mutagenesis experiment.

With a MinION flow cell, LevSeq can generate reads for a theoretical value of 2,500 96-well plates in a single flow cell, assuming 1,000 base pair lengths, as noted in previous nanopore sequencing methods^27^. Sequencing bias, increased sequencing length, and low-quality samples will reduce the number of useful sequences obtained. However, the reusability of the flow cell makes LevSeq a more economical option compared to Sanger and short-read next-generation sequencing. LevSeq is specifically designed to develop sequence-function datasets for research labs and as such the software has been designed for ease of installation and speed. A limitation of this is that the increased demultiplexing speed results in fewer reads assigned to each well compared with other methods^27,28^. For LevSeq we opted for this trade-off, which is beneficial when running experiments on a per lab basis where real-time data analysis is important for guiding the next step of an experiment. For industrial scale protein engineering procedures or in a sequencing facility, ParSEQ may be a more suitable pipeline^28^.

### Use case 2: Collecting sequence-function data to optimize variants using ML

To use *in silico* tools for protein optimization, one ideally starts with sequence-function data for a small subset of the studied system and these datasets serve as a foundation from which to make predictions and guide the engineering process. As one use case, LevSeq was used for active learning-assisted directed evolution (ALDE)^30^ by Yang et al. to sequence and analyze four 96-well plates of *Par*Pgb LQ variants from a 5-site combinatorial library. From the four plates, 216 unique variants without stop codons were identified and screened for activity and specificity for catalyzing the formation of the *cis* and *trans* cyclopropanation products of 4-methoxystyrene, the same reaction as in case 1. The sequence data from LevSeq and corresponding labels were used as initial training data for a batch Bayesian optimization algorithm, forming the baseline distribution to capture the effect on function of different amino acids at specific residues. This model was then used to suggest 96 sequences for testing; the researchers ordered exact genes for the 96 designed variants. Through three active learning loops the yield of non-native cyclopropanation reaction increased from 12% to 99%, with a 14:1 *cis*:*trans* selectivity ratio^30^. LevSeq enabled a critical step of collecting sequence-fitness datasets for model training in ML-assisted workflows. This foundation enhanced the effectiveness of subsequent rounds of ML guided directed evolution, leading to more successful outcomes.

In addition to collecting data for active learning, LevSeq can be used for experimental validation of suggested variants from various MLPE tools such as focused training MLDE (ftMLDE)^13^, cluster learning-assisted directed evolution (CLADE)^52^, and degenerate codon optimization for informed libraries (DeCOIL)^53^.

## Conclusion

LevSeq serves three primary functions for protein engineering: optimizing directed evolution workflows, gathering sequence information for specific MLPE projects, and consolidating sequence-function data for training future generalizable MLPE models. During exploration of the vast sequence-function space, experimentalists often encounter library bias; bias can lead to time and resources wasted on evaluating low-quality libraries. By providing sequence information prior to screening, LevSeq ensures that the gathered data are useful, regardless of whether the fitness results are positive, negative, or neutral.

In addition to its role in optimizing directed evolution workflows, the LevSeq pipeline can be used to generate high-quality datasets for MLPE. The cost-effective and efficient sequencing of variants from random mutagenesis studies using LevSeq helps overcome the bottleneck of limited data. Moreover, the scalability of LevSeq allows for the generation of datasets from a wide range of mutagenesis experiments, further expanding the scope of MLPE applications and facilitating advancements in protein engineering. By creating diverse and representative datasets that capture relevant sequence-function relationships, LevSeq can enable more robust and accurate models to be trained, ultimately leading to improved protein engineering and design.

## Supporting information

Complete supplemental information for LevSeq

## Acknowledgments

The authors thank Daniel Graves for initial primer design and Ravi Lai, Jason Yang, Kathleen Sicinski, Ziyang Qin, Julia Reisenbauer, Casey Ritts, and Jae Kennemeur for providing samples for testing and developing protocols. This material is based upon work supported by the U.S. Department of Energy, Office of Science, Office of Basic Energy Sciences, under Award Number DE-SC0022218. A.M. is Supported by the Schmidt Science Fellows, in partnership with the Rhodes Trust.

## Supporting Information Available

The following files are available free of charge.

- Supplementary materials for LevSeq: Rapid Generation of Sequence-Function Data for Directed Evolution and Machine Learning

