## Supplementary material for "LevSeq: Rapid Generation of Sequence-Function Data for Directed Evolution and Machine Learning": Complete supplemental information for LevSeq

### Oligonucleotide Design

#### Barcode Design

LevSeq uses 96 unique 24-nucleotide forward barcodes and 96 unique 24-nucleotide reverse barcodes. All barcodes have at least a hamming distance of three between each other (different by three substitutions). These barcodes were directly taken from the nanopore native barcoding kit and used with primers designed in the “Primer Design” section below.

#### Primer Design

The primers of LevSeq are specific to the cloning vectors that host the protein of interest (pET22b(+) in this protocol). Each full-length DNA sequence is captured by the combination of forward and reverse primers. These primers have the below general layout:

F: 5’ - XXXXXXXXXXXXXXXXXXXXXXXXATCTCGATCCCGCGAAATTAATACGACTCAC - 3’

R: 5’ – XXXXXXXXXXXXXXXXXXXXXXXXGCCCCAAGGGGTTATGCTAGTTATTGCTC – 3’

The 3’ region is a backbone-specific primer that binds to the cloning vector near the T7 and T7’ promotor sites. The 5’ region is the unique 24-nucleotide primer used for differentiating between plates (R) and wells (F).

### Supplemental Protocols

#### Ordering Barcode Linked Primers

The forward and reverse 24-nucleotide barcode sequences are given in Tables S1and S2; the full-length barcode linked forward and reverse primers for pET22b(+) are given in Tables S3 and S4. If using pET22B(+) as a cloning vector, the sequences listed in Tables S3 and S4 can be directly used for ordering primers at 100 µM concentrations from commercial suppliers. If using a different cloning vector, the user must verify that the primers in the “Primer Design” section would still amplify the desired region.

#### Preparation of LevSeq Barcode Primer Mixes

There are 96 unique forward barcode-linked primers, corresponding to each well of a 96-well plate; there are 96 reverse barcode-linked primers, corresponding to the ability to sequence 96 different plates as a unique reverse barcode is used for each plate. The barcode-linked primers are listed in the tables below (Tables S3 and S4); the 96 F primers and 96 R primers are both ordered in plate format at 100 µM concentration.

Each well sequenced using LevSeq is encoded by a unique combination of a forward (F) and a reverse (R) barcode. The reverse barcode would identify the sequence to a specific plate, and the forward barcode would identify the sequence to a specific well. Since these barcode-linked primers are compatible with any gene cloned in a specific cloning vector, it is convenient to keep a set of barcode-linked primer plates on hand.

We used the same eight barcode plates throughout this work, and they are named LevSeq01–LevSeq08. The exact barcodes used are given in Tables S5–12. The LevSeq software assumes the forward barcodes used for library preparation are laid out in the order given in Tables S5 and S12. To build the barcode plates depicted in Tables S5–12, we followed the below procedure:

1. A 10-fold dilution of the 100 µM forward barcode-linked primer plate from IDT was prepared by adding 10 µL of the forward primer stock to 90 µL dd$H_2O$ to a final concentration of 10 µM, keeping the well layout constant. Dilutions were performed in half skirted PCR plates (BIO-RAD HSP9601).
2. A 10-fold dilution of the reverse barcode-linked primer from IDT was prepared by adding 150 µL of the 100 µM reverse primer stock to 1350 µL dd$H_2O$ to a final concentration of 10 µM. Dilutions were performed in Eppendorf tubes.
3. To create plates for sequencing of forward/reverse primer mixes, 80 µL dd$H_2O$ were added to each well of each of the eight plates. Then, 10 µL diluted (10 µM) forward barcode plate was aliquoted to each well keeping the layout constant.
4. A unique reverse barcode was added to every well of each of the eight plates. To do so, 10 µL of diluted (10 µM) reverse barcode from step 2 was used. For instance, LevSeq01 has forward barcode 01 – 96 and reverse barcode 01, LevSeq02 has forward barcode 01 – 96 and reverse barcode 02, and so on. The final concentration of each barcode plate is 1 µM each for the forward and reverse primers (10X stock).
5. Eight of the LevSeq barcode plates (LevSeq 01 – 08) at 10X stock concentration were made in step 4. In order to make 1X reaction-ready barcode plates, the following dilution was performed: To another set of eight fully skirted PCR plates, 90 µL of dd$H_2O$ were added, followed by 10 µL of diluted LevSeq barcode stock plates (1 µM). The final concentration of each ready-to-use barcode plate is 0.1 µM for both forward and reverse barcode linked primers.
6. When not in use, the 10X stocks prepared in step 4 were stored at -20 °C, while the barcode plates were stored at 4 °C. The 100 µM, 10 µM, and 1 µM barcode plates can be stored for long periods of time.

#### LevSeq Library Preparation and Sequencing

The steps below can be followed to complete a LevSeq sample preparation ready for loading onto an Oxford Nanopore flow cell (FLO-MIN114). Note that when designing a new set of barcode-linked backbone-specific primers, it is recommended to test the primers and PCR protocol using a few wells before deploying them for plate-scale reactions. The library protocol below provides part numbers that are for the materials and reagents used while developing this protocol, reagents from other providers should work as well.

1. Prepare a PCR master mix for the number of plates to be sequenced according to the table below.

| Component | Amount per plate (µL) |
|---|---|
| Thermopol Buffer (NEB B9004L) | 144 |
| 10 mM dNTPs (NEB N0447) | 28.8 |
| Taq Polymerase (NEB M0267) | 7.2 |
| Mol-Bio Grade DMSO (mp 194819) | 57.6 |
| ddH2O | 770 |

2. Add 7 µL of master mix to each well of as many half-skirted PCR plates (USA scientific 1402-9700) as will be sequenced. These are referred to as “PCR plates”.
3. Stamp 2 µL of the 1-µM barcode-linked primer mix from the barcode plates into the PCR plates.
    a. “Stamp” means “apply to all wells, keeping the plate layout consistent”.
4. Stamp 1 µL of overnight culture from each plate to be sequenced into the PCR plates. **Record which barcode plate was used with which PCR plate.**
5. Complete a PCR using the below thermal cycler program. This colony PCR amplifies the entire gene of interest from the template DNA contained in the cell culture.

| Step | Temperature (°C) | Time |
|---|---|---|
| 1 | 95 | 5 min |
| 2 | 95 | 20 s |
| 3 | TD 68 -> 63.5 | 20 s |
| 4 | 68 | 1 min per kb |
| 5 | Return to 2, 9x | |
| 6 | 95 | 20 s |
| 7 | 68 | 1 min per kb |
| 8 | Return to 6, 24x | |
| 9 | 68 | 5 min |
| 10 | 4 | Hold |

    a. “TD” in step 3 stands for “touchdown”, a touchdown step decreases the temperature each cycle. The TD in the above PCR starts at 68 °C and drops to 63.5 °C by the end, decreasing by 0.5 °C per cycle.
    b. Extension time in steps 4 and 7 are recommended to be minimally 1 minute per kb of gene, a 2-min extension time was used for genes below 1 kb during development of this protocol.
6. While the PCR is running, prepare a 1% agarose gel with SYBR gold added (Thermo Fisher Scientific, S11494).
7. Once the PCR is completed, for each plate, pool 5 µL of each reaction into an Eppendorf tube, for a total of 480 µL of pooled PCR products per plate. Pooling will leave you with as many Eppendorf tubes as you have plates.
    a. Note: Pooling can be performed using a 12-channel multichannel pipette. 1) First, transfer 5 µL of reactions from each row in the PCR plate (to be sequenced) into a new row of a 96-well PCR plate. Since there are eight rows in the original plate, each well of the row in the new 96-well PCR plate will have 40 µL of pooled PCR products. 2) Transfer 30 µL from each well in the single row of pooled reactions using a single-channel pipette to a microcentrifuge tube. Since there are 12 columns in the new 96-well PCR plate, the microcentrifuge tube should have 360 µL of pooled PCR products that contain PCR reactions of every well of the PCR plate (to be sequenced).
    b. Alternatively, a liquid handling robot can be used for this task. **It is important that 5 µL from each well of each PCR plate is combined into one microcentrifuge tube to improve the evenness of sequencing coverage. There will now be an equal number of microcentrifuge tubes to as plates being sequenced.**
8. For each tube made in step 7, take 100 µL of pooled PCR reactions and add it to 20 µL 6x loading dye (NEB B7025S) in a microcentrifuge tube. The remaining pooled reaction can be stored at -20 °C for future use.
9. Load the contents of each tube made in step 9 into the agarose gel prepared in step 7. Each tube should have separated lanes. Load a 10 µL of 1 kb ladder in the flanking lanes.
10. Run the agarose gel at 120 V until the bands have sufficiently migrated. Reference the ladder to identify the PCR products which should be the size of the gene plus about 100 bp. The extra bases come from the barcodes and the region between the primer attachment sites and the open reading frame.
11. Gel extract the desired bands. Samples from separated PCR reaction plates should have their own microcentrifuge tube to hold the extracted gels. Commercial kits such as the Zymoclean Gel DNA Recovery Kit (Zymo Research, D4001) are typically used for this step. Elution should be performed using 10 µL of $ddH_2O$ rather than elution buffer.
12. After gel extraction, measure the DNA concentration of each gel-extracted PCR product from each plate. We use a GE NanoVue Plus to measure DNA concentration in ng/µL unit.
13. Combine the gel-extracted PCR products from each plate in equimolar concentrations. In general, 200 ng of pooled DNA sample per 1-kb gene are sufficient for any sample preparation. For experiments that use the

same length template, the weight is directly proportional to molarity. Hence the same amount of DNA in nanograms from each plate can be combined into one microcentrifuge tube. For example, if plate one after gel extraction has a measured concentration of 20 ng/μL and plate two has a concentration of 10 ng/μL, then to make 20 μL of 10 ng/μL final sample, 5 μL of the first plate and 10 μL of the second plate were added to 5 μL of ddH2O to make the final sample.

14. After step 13, there should be one single tube of cleaned, normalized DNA consisting of all genes from all plates to be sequenced. Depending on the Oxford Nanopore sample preparation protocol, this sample is adjusted to the desired concentration before sample preparation for sequencing.
15. A MinION flow cell with the LSK114 ligation sequencing kit was used while developing this protocol, which recommends 200 fmol of amplicon DNA for sample preparation. This translates to 120 ng for a 1-kb gene.
16. From the concentration-normalized sample prepared in step 13, transfer the equivalent volume of 120 ng of pooled DNA into a new microcentrifuge tube. This pooled DNA sample (120 ng) will be carried forward for Oxford Nanopore sequencing sample preparation. Protocols are linked here (https://community.nanoporetech.com/docs/prepare/library_prep_protocols/genomic-dna-by-ligation-sqk-lsk114/v/gde_9161_v114_revu_29jun2022/dna-repair-and-end-prep?devices=minion).
17. Once the sample is loaded onto the flow cell for sequencing, we recommend choosing the super accurate basecalling option to ensure retrieval of the highest quality sequences while setting up the sequencing run. It is recommended to stop the flow cell after the basecalled bases reach a data volume of 0.02 Gb per plate to be sequenced, so an experiment with 20 plates would be stopped after collecting 0.4 Gb of basecalled bases.

#### LevSeq Data Analysis

If basecalling is enabled, a “fastq_pass” folder with the basecalled sequencing results will be returned at the designated folder location. This folder location is given at the end of the sequencing run set up and is usually where the Minknow software is installed in the system. (In our case, data can be accessed from “/var/lib/minknow/data/name_of_experiment”). Once sequencing is complete, LevSeq software can be used to process results (in the format of .fastq.gz files) and assign variants to their original wells. Detailed instructions on how to use the LevSeq software and interpret its outputs are provided on the LevSeq Wiki.

### Supplemental Figures

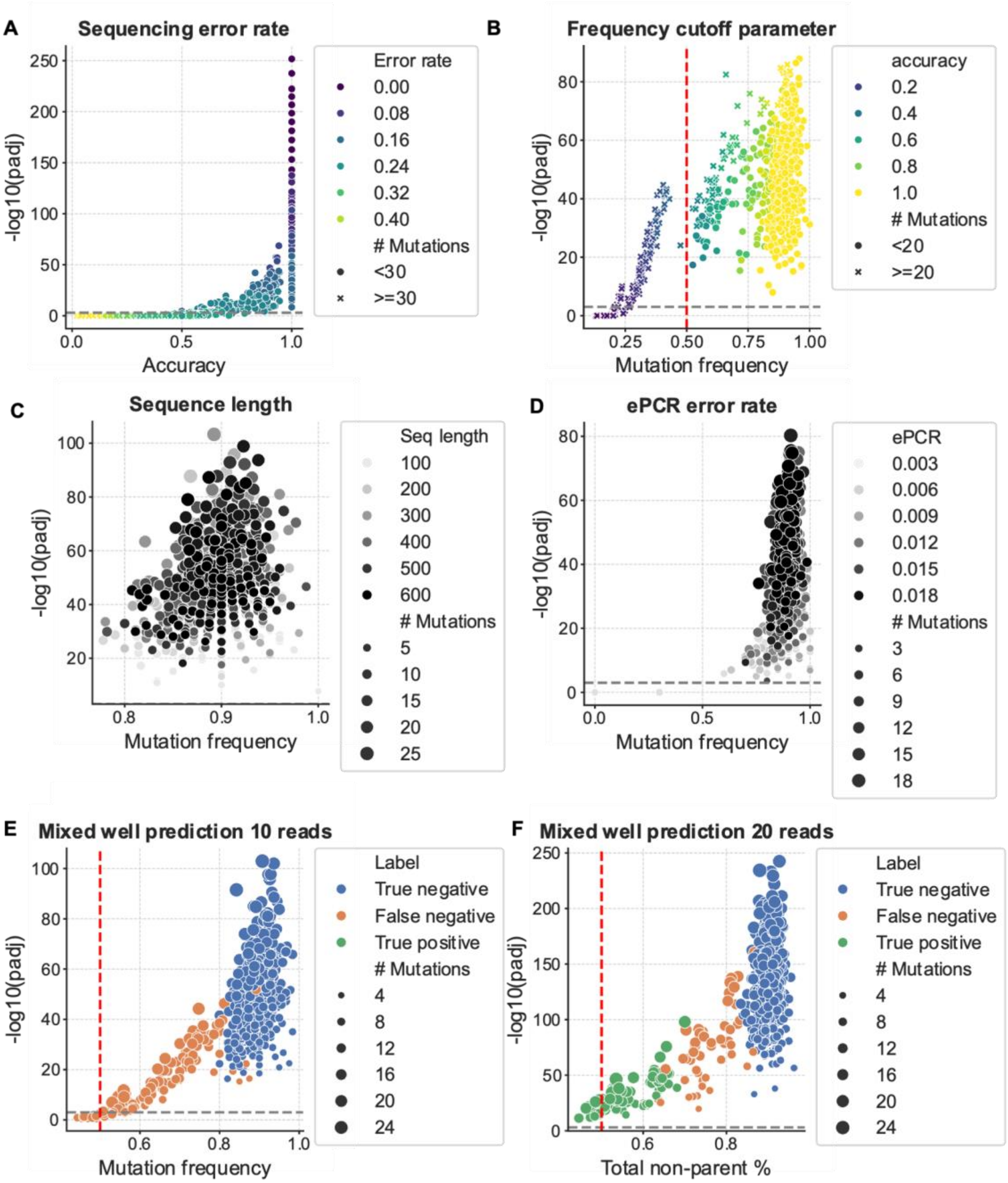

Figure S1: epPCR simulation study results. A. Nanopore sequencing error was varied from 0 to 50% at a step size of 5, holding the ePCR mutation rate constant at 2%, read depth of 10, and a frequency cutoff of 50%, which is default in LevSeq. B. Mutation frequency is a parameter that defines when a mutation is tested to be a true mutation from PCR (significant) rather than a sequencing error. The default is 50%, this means there must be 50% non-parent mutations within the position for significance to be tested, this reduces false positive rates and increases the accuracy of variant calling. C. Sequence length was varied while holding the other parameters constant, no trend is observed showing sequence length does not affect the identification of correct sequence, so gene length does not affect our software's variant calling ability. D. epPCR rate was varied up to an error rate of 2%, and no adverse effects were observed on the variant calling accuracy. E. Mixed wells were simulated by combining wells (doping in one sequence at a certain percentage), for these tests it was up to 50%, holding the other parameters constant. For 10 reads there is a high false negative rate, with mixed wells unable to be accurately predicted at the low sequence rate. F. Simulated as in E, except with 20 reads, the false negative rate is reduced and true positives are able to be detected, yielding an accuracy of 0.91, precision of 1.0, and recall of 0.59.

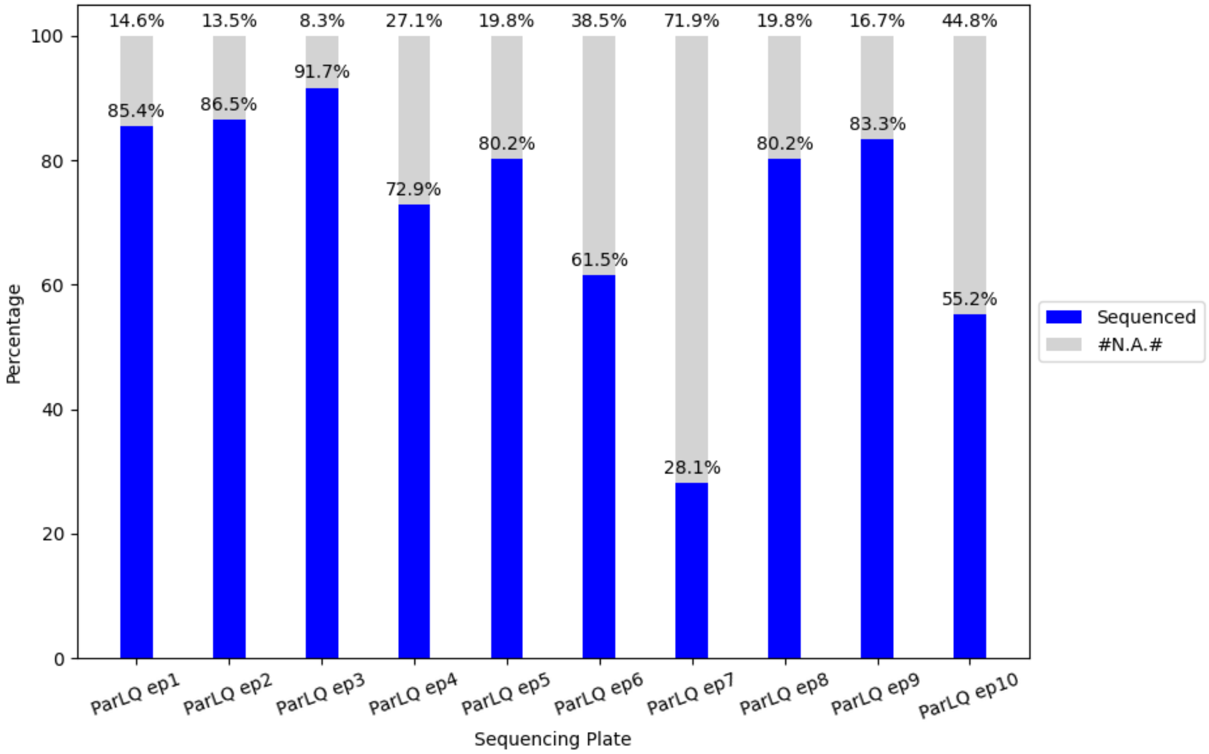

Figure S2: ParLQ (~200 amino acids) error prone library sequencing percentage by experiment plate. Plate 6, 7, and 10 had below 60% variants sequenced while the other seven plates have above 70% variants sequenced. This sequencing run was performed without optimizing PCR procedure.

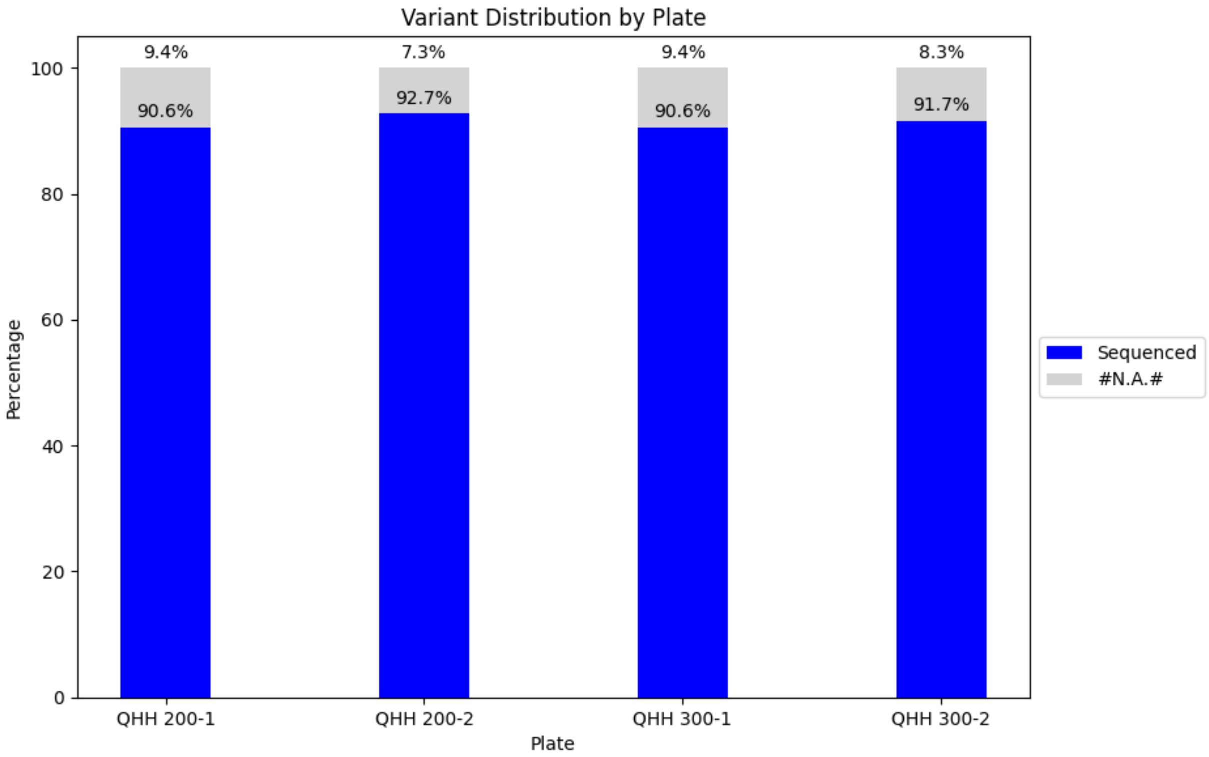

Figure S3: QHH (~500 amino acids) error-prone library sequencing percentage by experiment plate. This sequencing run is performed with updated PCR procedure.

### Supplemental Tables

### Barcode and Primer Sequences

**Table S1.** LevSeq forward barcode sequences used in this work. NB indicates forward barcodes.

| Well Position | Name | Barcode |
|---|---|---|
| A1 | NB01 | CACAAAGACACCGACAACTTTCTT |
| A2 | NB02 | ACAGACGACTACAAACGGAATCGA |
| A3 | NB03 | CCTGGTAACTGGGACACAAGACTC |
| A4 | NB04 | TAGGGAAACACGATAGAATCCGAA |
| A5 | NB05 | AAGGTTACACAAACCCTGGACAAG |
| A6 | NB06 | GACTACTTTCTGCCTTTGCGAGAA |
| A7 | NB07 | AAGGATTCATTCCCACGGTAACAC |
| A8 | NB08 | ACGTAACTTGGTTTGTTCCCTGAA |
| A9 | NB09 | AACCAAGACTCGCTGTGCCTAGTT |
| A10 | NB10 | GAGAGGACAAAGGTTTCAACGCTT |
| A11 | NB11 | TCCATTCCCTCCGATAGATGAAAC |
| A12 | NB12 | TCCGATTCTGCTTCTTTCTACCTG |
| B1 | NB13 | AGAACGACTTCCATACTCGTGTGA |
| B2 | NB14 | AACGAGTCTCTTGGGACCCATAGA |
| B3 | NB15 | AGGTCTACCTCGCTAACACCACTG |
| B4 | NB16 | CGTCAACTGACAGTGGTTCGTACT |
| B5 | NB17 | ACCCTCCAGGAAAGTACCTCTGAT |
| B6 | NB18 | CCAAACCCAACAACCTAGATAGGC |
| B7 | NB19 | GTTCCTCGTGCAGTGTCAAGAGAT |
| B8 | NB20 | TTGCGTCCTGTTACGAGAACTCAT |
| B9 | NB21 | GAGCCTCTCATTGTCCGTTCTCTA |
| B10 | NB22 | ACCACTGCCATGTATCAAAGTACG |
| B11 | NB23 | CTTACTACCCAGTGAACCTCCTCG |
| B12 | NB24 | GCATAGTTCTGCATGATGGGTTAG |
| C1 | NB25 | GTAAGTTGGGTATGCAACGCAATG |
| C2 | NB26 | CATACAGCGACTACGCATTCTCAT |
| C3 | NB27 | CGACGGTTAGATTCACCTCTTACA |
| C4 | NB28 | TGAAACCTAAGAAGGCACCGTATC |
| C5 | NB29 | CTAGACACCTTGGGTTGACAGACC |
| C6 | NB30 | TCAGTGAGGATCTACTTCGACCCA |
| C7 | NB31 | TGCGTACAGCAATCAGTTACATTG |
| C8 | NB32 | CCAGTAGAAGTCCGACAACGTCAT |
| C9 | NB33 | CAGACTTGGTACGGTTGGGTAACT |

| | | |
|---|---|---|
| C10 | NB34 | GGACGAAGAACTCAAGTCAAAGGC |
| C11 | NB35 | CTACTTACGAAGCTGAGGGACTGC |
| C12 | NB36 | ATGTCCCAGTTAGAGGAGGAAACA |
| D1 | NB37 | GCTTGCGATTGATGCTTAGTATCA |
| D2 | NB38 | ACCACAGGAGGACGATACAGAGAA |
| D3 | NB39 | CCACAGTGTCAACTAGAGCCTCTC |
| D4 | NB40 | TAGTTTGGATGACCAAGGATAGCC |
| D5 | NB41 | GGAGTTCGTCCAGAGAAGTACACG |
| D6 | NB42 | CTACGTGTAAGGCATACCTGCCAG |
| D7 | NB43 | CTTTCGTTGTTGACTCGACGGTAG |
| D8 | NB44 | AGTAGAAAGGGTTCCTTCCCACTC |
| D9 | NB45 | GATCCAACAGAGATGCCTTCAGTG |
| D10 | NB46 | GCTGTGTTCCACTTCATTCTCCTG |
| D11 | NB47 | GTGCAACTTTCCCACAGGTAGTTC |
| D12 | NB48 | CATCTGGAACGTGGTACACCTGTA |
| E1 | NB49 | ACTGGTGCAGCTTTGAACATCTAG |
| E2 | NB50 | ATGGACTTTGGTAACTTCCTGCGT |
| E3 | NB51 | GTTGAATGAGCCTACTGGGTCCTC |
| E4 | NB52 | TGAGAGACAAGATTGTTCGTGGAC |
| E5 | NB53 | AGATTCAGACCGTCTCATGCAAAG |
| E6 | NB54 | CAAGAGCTTTGACTAAGGAGCATG |
| E7 | NB55 | TGGAAGATGAGACCCTGATCTACG |
| E8 | NB56 | TCACTACTCAACAGGTGGCATGAA |
| E9 | NB57 | GCTAGGTCAATCTCCTTCGGAAGT |
| E10 | NB58 | CAGGTTACTCCTCCGTGAGTCTGA |
| E11 | NB59 | TCAATCAAGAAGGGAAAGCAAGGT |
| E12 | NB60 | CATGTTCAACCAAGGCTTCTATGG |
| F1 | NB61 | AGAGGGTACTATGTGCCTCAGCAC |
| F2 | NB62 | CACCCACACTTACTTCAGGACGTA |
| F3 | NB63 | TTCTGAAGTTCCTGGGTCTTGAAC |
| F4 | NB64 | GACAGACACCGTTCATCGACTTTC |
| F5 | NB65 | TTCTCAGTCTTCCTCCAGACAAGG |
| F6 | NB66 | CCGATCCTTGTGGCTTCTAACTTC |
| F7 | NB67 | GTTTGTCATACTCGTGTGCTCACC |
| F8 | NB68 | GAATCTAAGCAAACACGAAGGTGG |
| F9 | NB69 | TACAGTCCGAGCCTCATGTGATCT |
| F10 | NB70 | ACCGAGATCCTACGAATGGAGTGT |
| F11 | NB71 | CCTGGGAGCATCAGGTAGTAACAG |
| F12 | NB72 | TAGCTGACTGTCTTCCATACCGAC |

| | | |
|---|---|---|
| G1 | NB73 | AAGAAACAGGATGACAGAACCCTC |
| G2 | NB74 | TACAAGCATCCCAACACTTCCACT |
| G3 | NB75 | GACCATTGTGATGAACCCTGTTGT |
| G4 | NB76 | ATGCTTGTTACATCAACCCTGGAC |
| G5 | NB77 | CGACCTGTTTCTCAGGGATACAAC |
| G6 | NB78 | AACAACCGAACCTTTGAATCAGAA |
| G7 | NB79 | TCTCGGAGATAGTTCTCACTGCTG |
| G8 | NB80 | CGGATGAACATAGGATAGCGATTC |
| G9 | NB81 | CCTCATCTTGTGAAGTTGTTTCGG |
| G10 | NB82 | ACGGTATGTCGAGTTCCAGGACTA |
| G11 | NB83 | TGGCTTGATCTAGGTAAGGTCGAA |
| G12 | NB84 | GTAGTGGACCTAGAACCTGTGCCA |
| H1 | NB85 | AACGGAGGAGTTAGTTGGATGATC |
| H2 | NB86 | AGGTGATCCCAACAAGCGTAAGTA |
| H3 | NB87 | TACATGCTCCTGTTGTTAGGGAGG |
| H4 | NB88 | TCTTCTACTACCGATCCGAAGCAG |
| H5 | NB89 | ACAGCATCAATGTTTGGCTAGTTG |
| H6 | NB90 | GATGTAGAGGGTACGGTTTGAGGC |
| H7 | NB91 | GGCTCCATAGGAACTCACGCTACT |
| H8 | NB92 | TTGTGAGTGGAAAGATACAGGACC |
| H9 | NB93 | AGTTTCCATCACTTCAGACTTGGG |
| H10 | NB94 | GATTGTCCTCAAACTGCCACCTAC |
| H11 | NB95 | CCTGTCTGGAAGAAGAATGGACTT |
| H12 | NB96 | CTGAACGGTCATAGAGTCCACCAT |

**Table S2.** LevSeqreverse barcode sequences used in this work. RB stands for reverse barcodes.

| Well Position | Name | Barcode |
|---|---|---|
| A1 | RB01 | AAGAAAGTTGTCGGTGTCTTTGTG |
| A2 | RB02 | TCGATTCCGTTTGTAGTCGTCTGT |
| A3 | RB03 | GAGTCTTGTGTCCCAGTTACCAGG |
| A4 | RB04 | TTCGGATTCTATCGTGTTTCCCTA |
| A5 | RB05 | CTTGTCCAGGGTTTGTGTAACCTT |
| A6 | RB06 | TTCTCGCAAAGGCAGAAAGTAGTC |
| A7 | RB07 | GTGTTACCGTGGGAATGAATCCTT |
| A8 | RB08 | TTCAGGGAACAAACCAAGTTACGT |
| A9 | RB09 | AACTAGGCACAGCGAGTCTTGGTT |
| A10 | RB10 | AAGCGTTGAAACCTTTGTCCTCTC |
| A11 | RB11 | GTTTCATCTATCGGAGGGAATGGA |
| A12 | RB12 | CAGGTAGAAAGAAGCAGAATCGGA |

| | | |
|---|---|---|
| B1 | RB13 | AGAACGACTTCCATACTCGTGTGA |
| B2 | RB14 | AACGAGTCTCTTGGGACCCATAGA |
| B3 | RB15 | AGGTCTACCTCGCTAACACCACTG |
| B4 | RB16 | CGTCAACTGACAGTGGTTCGTACT |
| B5 | RB17 | ACCCTCCAGGAAAGTACCTCTGAT |
| B6 | RB18 | CCAAACCCAACAACCTAGATAGGC |
| B7 | RB19 | GTTCCTCGTGCAGTGTCAAGAGAT |
| B8 | RB20 | TTGCGTCCTGTTACGAGAACTCAT |
| B9 | RB21 | GAGCCTCTCATTGTCCGTTCTCTA |
| B10 | RB22 | ACCACTGCCATGTATCAAAGTACG |
| B11 | RB23 | CTTACTACCCAGTGAACCTCCTCG |
| B12 | RB24 | GCATAGTTCTGCATGATGGGTTAG |
| C1 | RB25 | GTAAGTTGGGTATGCAACGCAATG |
| C2 | RB26 | CATACAGCGACTACGCATTCTCAT |
| C3 | RB27 | CGACGGTTAGATTCACCTCTTACA |
| C4 | RB28 | TGAAACCTAAGAAGGCACCGTATC |
| C5 | RB29 | CTAGACACCTTGGGTTGACAGACC |
| C6 | RB30 | TCAGTGAGGATCTACTTCGACCCA |
| C7 | RB31 | TGCGTACAGCAATCAGTTACATTG |
| C8 | RB32 | CCAGTAGAAGTCCGACAACGTCAT |
| C9 | RB33 | CAGACTTGGTACGGTTGGGTAACT |
| C10 | RB34 | GGACGAAGAACTCAAGTCAAAGGC |
| C11 | RB35 | CTACTTACGAAGCTGAGGGACTGC |
| C12 | RB36 | ATGTCCCAGTTAGAGGAGGAAACA |
| D1 | RB37 | GCTTGCGATTGATGCTTAGTATCA |
| D2 | RB38 | ACCACAGGAGGACGATACAGAGAA |
| D3 | RB39 | CCACAGTGTCAACTAGAGCCTCTC |
| D4 | RB40 | TAGTTTGGATGACCAAGGATAGCC |
| D5 | RB41 | GGAGTTCGTCCAGAGAAGTACACG |
| D6 | RB42 | CTACGTGTAAGGCATACCTGCCAG |
| D7 | RB43 | CTTTCGTTGTTGACTCGACGGTAG |
| D8 | RB44 | AGTAGAAAGGGTTCCTTCCCACTC |
| D9 | RB45 | GATCCAACAGAGATGCCTTCAGTG |
| D10 | RB46 | GCTGTGTTCCACTTCATTCTCCTG |
| D11 | RB47 | GTGCAACTTTCCCACAGGTAGTTC |
| D12 | RB48 | CATCTGGAACGTGGTACACCTGTA |
| E1 | RB49 | ACTGGTGCAGCTTTGAACATCTAG |
| E2 | RB50 | ATGGACTTTGGTAACTTCCTGCGT |
| E3 | RB51 | GTTGAATGAGCCTACTGGGTCCTC |

| | | |
|---|---|---|
| E4 | RB52 | TGAGAGACAAGATTGTTCGTGGAC |
| E5 | RB53 | AGATTCAGACCGTCTCATGCAAAG |
| E6 | RB54 | CAAGAGCTTTGACTAAGGAGCATG |
| E7 | RB55 | TGGAAGATGAGACCCTGATCTACG |
| E8 | RB56 | TCACTACTCAACAGGTGGCATGAA |
| E9 | RB57 | GCTAGGTCAATCTCCTTCGGAAGT |
| E10 | RB58 | CAGGTTACTCCTCCGTGAGTCTGA |
| E11 | RB59 | TCAATCAAGAAGGGAAAGCAAGGT |
| E12 | RB60 | CATGTTCAACCAAGGCTTCTATGG |
| F1 | RB61 | AGAGGGTACTATGTGCCTCAGCAC |
| F2 | RB62 | CACCCACACTTACTTCAGGACGTA |
| F3 | RB63 | TTCTGAAGTTCCTGGGTCTTGAAC |
| F4 | RB64 | GACAGACACCGTTCATCGACTTTC |
| F5 | RB65 | TTCTCAGTCTTCCTCCAGACAAGG |
| F6 | RB66 | CCGATCCTTGTGGCTTCTAACTTC |
| F7 | RB67 | GTTTGTCATACTCGTGTGCTCACC |
| F8 | RB68 | GAATCTAAGCAAACACGAAGGTGG |
| F9 | RB69 | TACAGTCCGAGCCTCATGTGATCT |
| F10 | RB70 | ACCGAGATCCTACGAATGGAGTGT |
| F11 | RB71 | CCTGGGAGCATCAGGTAGTAACAG |
| F12 | RB72 | TAGCTGACTGTCTTCCATACCGAC |
| G1 | RB73 | AAGAAACAGGATGACAGAACCCTC |
| G2 | RB74 | TACAAGCATCCCAACACTTCCACT |
| G3 | RB75 | GACCATTGTGATGAACCCTGTTGT |
| G4 | RB76 | ATGCTTGTTACATCAACCCTGGAC |
| G5 | RB77 | CGACCTGTTTCTCAGGGATACAAC |
| G6 | RB78 | AACAACCGAACCTTTGAATCAGAA |
| G7 | RB79 | TCTCGGAGATAGTTCTCACTGCTG |
| G8 | RB80 | CGGATGAACATAGGATAGCGATTC |
| G9 | RB81 | CCTCATCTTGTGAAGTTGTTTCGG |
| G10 | RB82 | ACGGTATGTCGAGTTCCAGGACTA |
| G11 | RB83 | TGGCTTGATCTAGGTAAGGTCGAA |
| G12 | RB84 | GTAGTGGACCTAGAACCTGTGCCA |
| H1 | RB85 | AACGGAGGAGTTAGTTGGATGATC |
| H2 | RB86 | AGGTGATCCCAACAAGCGTAAGTA |
| H3 | RB87 | TACATGCTCCTGTTGTTAGGGAGG |
| H4 | RB88 | TCTTCTACTACCGATCCGAAGCAG |
| H5 | RB89 | ACAGCATCAATGTTTGGCTAGTTG |
| H6 | RB90 | GATGTAGAGGGTACGGTTTGAGGC |

| | | |
|---|---|---|
| H7 | RB91 | GGCTCCATAGGAACTCACGCTACT |
| H8 | RB92 | TTGTGAGTGGAAAGATACAGGACC |
| H9 | RB93 | AGTTTCCATCACTTCAGACTTGGG |
| H10 | RB94 | GATTGTCCTCAAACTGCCACCTAC |
| H11 | RB95 | CCTGTCTGGAAGAAGAATGGACTT |
| H12 | RB96 | CTGAACGGTCATAGAGTCCACCAT |

**Table S3.** Full-length LevSeq forward barcode-linked primer sequences used in this work. The primer-linked barcodes below are directly ordered from IDT.

| Well Position | Name | Barcode linked primer sequence |
|---|---|---|
| A1 | NB01 | CACAAAGACACCGACAACTTTCTTATCTCGATCCCGCGAAATTAATACGACTCAC |
| A2 | NB02 | ACAGACGACTACAAACGGAATCGAATCTCGATCCCGCGAAATTAATACGACTCAC |
| A3 | NB03 | CCTGGTAACTGGGACACAAGACTCATCTCGATCCCGCGAAATTAATACGACTCAC |
| A4 | NB04 | TAGGGAAACACGATAGAATCCGAAATCTCGATCCCGCGAAATTAATACGACTCAC |
| A5 | NB05 | AAGGTTACACAAACCCTGGACAAGATCTCGATCCCGCGAAATTAATACGACTCAC |
| A6 | NB06 | GACTACTTTCTGCCTTTGCGAGAAATCTCGATCCCGCGAAATTAATACGACTCAC |
| A7 | NB07 | AAGGATTCATTCCCACGGTAACACATCTCGATCCCGCGAAATTAATACGACTCAC |
| A8 | NB08 | ACGTAACTTGGTTTGTTCCCTGAAATCTCGATCCCGCGAAATTAATACGACTCAC |
| A9 | NB09 | AACCAAGACTCGCTGTGCCTAGTTATCTCGATCCCGCGAAATTAATACGACTCAC |
| A10 | NB10 | GAGAGGACAAAGGTTTCAACGCTTATCTCGATCCCGCGAAATTAATACGACTCAC |
| A11 | NB11 | TCCATTCCCTCCGATAGATGAAACATCTCGATCCCGCGAAATTAATACGACTCAC |
| A12 | NB12 | TCCGATTCTGCTTCTTTCTACCTGATCTCGATCCCGCGAAATTAATACGACTCAC |
| B1 | NB13 | AGAACGACTTCCATACTCGTGTGAATCTCGATCCCGCGAAATTAATACGACTCAC |
| B2 | NB14 | AACGAGTCTCTTGGGACCCATAGAATCTCGATCCCGCGAAATTAATACGACTCAC |
| B3 | NB15 | AGGTCTACCTCGCTAACACCACTGATCTCGATCCCGCGAAATTAATACGACTCAC |
| B4 | NB16 | CGTCAACTGACAGTGGTTCGTACTATCTCGATCCCGCGAAATTAATACGACTCAC |
| B5 | NB17 | ACCCTCCAGGAAAGTACCTCTGATATCTCGATCCCGCGAAATTAATACGACTCAC |
| B6 | NB18 | CCAAACCCAACAACCTAGATAGGCATCTCGATCCCGCGAAATTAATACGACTCAC |
| B7 | NB19 | GTTCCTCGTGCAGTGTCAAGAGATATCTCGATCCCGCGAAATTAATACGACTCAC |
| B8 | NB20 | TTGCGTCCTGTTACGAGAACTCATATCTCGATCCCGCGAAATTAATACGACTCAC |
| B9 | NB21 | GAGCCTCTCATTGTCCGTTCTCTAATCTCGATCCCGCGAAATTAATACGACTCAC |
| B10 | NB22 | ACCACTGCCATGTATCAAAGTACGATCTCGATCCCGCGAAATTAATACGACTCAC |
| B11 | NB23 | CTTACTACCCAGTGAACCTCCTCGATCTCGATCCCGCGAAATTAATACGACTCAC |
| B12 | NB24 | GCATAGTTCTGCATGATGGGTTAGATCTCGATCCCGCGAAATTAATACGACTCAC |
| C1 | NB25 | GTAAGTTGGGTATGCAACGCAATGATCTCGATCCCGCGAAATTAATACGACTCAC |
| C2 | NB26 | CATACAGCGACTACGCATTCTCATATCTCGATCCCGCGAAATTAATACGACTCAC |
| C3 | NB27 | CGACGGTTAGATTCACCTCTTACAATCTCGATCCCGCGAAATTAATACGACTCAC |
| C4 | NB28 | TGAAACCTAAGAAGGCACCGTATCATCTCGATCCCGCGAAATTAATACGACTCAC |
| C5 | NB29 | CTAGACACCTTGGGTTGACAGACCATCTCGATCCCGCGAAATTAATACGACTCAC |
| C6 | NB30 | TCAGTGAGGATCTACTTCGACCCAATCTCGATCCCGCGAAATTAATACGACTCAC |
| C7 | NB31 | TGCGTACAGCAATCAGTTACATTGATCTCGATCCCGCGAAATTAATACGACTCAC |
| C8 | NB32 | CCAGTAGAAGTCCGACAACGTCATATCTCGATCCCGCGAAATTAATACGACTCAC |
| C9 | NB33 | CAGACTTGGTACGGTTGGGTAACTATCTCGATCCCGCGAAATTAATACGACTCAC |
| C10 | NB34 | GGACGAAGAACTCAAGTCAAAGGCATCTCGATCCCGCGAAATTAATACGACTCAC |
| C11 | NB35 | CTACTTACGAAGCTGAGGGACTGCATCTCGATCCCGCGAAATTAATACGACTCAC |
| C12 | NB36 | ATGTCCCAGTTAGAGGAGGAAACAATCTCGATCCCGCGAAATTAATACGACTCAC |

| | | |
|---|---|---|
| D1 | NB37 | GCTTGCGATTGATGCTTAGTATCAATCTCGATCCCGCGAAATTAATACGACTCAC |
| D2 | NB38 | ACCACAGGAGGACGATACAGAGAAATCTCGATCCCGCGAAATTAATACGACTCAC |
| D3 | NB39 | CCACAGTGTCAACTAGAGCCTCTCATCTCGATCCCGCGAAATTAATACGACTCAC |
| D4 | NB40 | TAGTTTGGATGACCAAGGATAGCCATCTCGATCCCGCGAAATTAATACGACTCAC |
| D5 | NB41 | GGAGTTCGTCCAGAGAAGTACACGATCTCGATCCCGCGAAATTAATACGACTCAC |
| D6 | NB42 | CTACGTGTAAGGCATACCTGCCAGATCTCGATCCCGCGAAATTAATACGACTCAC |
| D7 | NB43 | CTTTCGTTGTTGACTCGACGGTAGATCTCGATCCCGCGAAATTAATACGACTCAC |
| D8 | NB44 | AGTAGAAAGGGTTCCTTCCCACTCATCTCGATCCCGCGAAATTAATACGACTCAC |
| D9 | NB45 | GATCCAACAGAGATGCCTTCAGTGATCTCGATCCCGCGAAATTAATACGACTCAC |
| D10 | NB46 | GCTGTGTTCCACTTCATTCTCCTGATCTCGATCCCGCGAAATTAATACGACTCAC |
| D11 | NB47 | GTGCAACTTTCCCACAGGTAGTTCATCTCGATCCCGCGAAATTAATACGACTCAC |
| D12 | NB48 | CATCTGGAACGTGGTACACCTGTAATCTCGATCCCGCGAAATTAATACGACTCAC |
| E1 | NB49 | ACTGGTGCAGCTTTGAACATCTAGATCTCGATCCCGCGAAATTAATACGACTCAC |
| E2 | NB50 | ATGGACTTTGGTAACTTCCTGCGTATCTCGATCCCGCGAAATTAATACGACTCAC |
| E3 | NB51 | GTTGAATGAGCCTACTGGGTCCTCATCTCGATCCCGCGAAATTAATACGACTCAC |
| E4 | NB52 | TGAGAGACAAGATTGTTCGTGGACATCTCGATCCCGCGAAATTAATACGACTCAC |
| E5 | NB53 | AGATTCAGACCGTCTCATGCAAAGATCTCGATCCCGCGAAATTAATACGACTCAC |
| E6 | NB54 | CAAGAGCTTTGACTAAGGAGCATGATCTCGATCCCGCGAAATTAATACGACTCAC |
| E7 | NB55 | TGGAAGATGAGACCCTGATCTACGATCTCGATCCCGCGAAATTAATACGACTCAC |
| E8 | NB56 | TCACTACTCAACAGGTGGCATGAAATCTCGATCCCGCGAAATTAATACGACTCAC |
| E9 | NB57 | GCTAGGTCAATCTCCTTCGGAAGTATCTCGATCCCGCGAAATTAATACGACTCAC |
| E10 | NB58 | CAGGTTACTCCTCCGTGAGTCTGAATCTCGATCCCGCGAAATTAATACGACTCAC |
| E11 | NB59 | TCAATCAAGAAGGGAAAGCAAGGTATCTCGATCCCGCGAAATTAATACGACTCAC |
| E12 | NB60 | CATGTTCAACCAAGGCTTCTATGGATCTCGATCCCGCGAAATTAATACGACTCAC |
| F1 | NB61 | AGAGGGTACTATGTGCCTCAGCACATCTCGATCCCGCGAAATTAATACGACTCAC |
| F2 | NB62 | CACCCACACTTACTTCAGGACGTAATCTCGATCCCGCGAAATTAATACGACTCAC |
| F3 | NB63 | TTCTGAAGTTCCTGGGTCTTGAACATCTCGATCCCGCGAAATTAATACGACTCAC |
| F4 | NB64 | GACAGACACCGTTCATCGACTTTCATCTCGATCCCGCGAAATTAATACGACTCAC |
| F5 | NB65 | TTCTCAGTCTTCCTCCAGACAAGGATCTCGATCCCGCGAAATTAATACGACTCAC |
| F6 | NB66 | CCGATCCTTGTGGCTTCTAACTTCATCTCGATCCCGCGAAATTAATACGACTCAC |
| F7 | NB67 | GTTTGTCATACTCGTGTGCTCACCATCTCGATCCCGCGAAATTAATACGACTCAC |
| F8 | NB68 | GAATCTAAGCAAACACGAAGGTGGATCTCGATCCCGCGAAATTAATACGACTCAC |
| F9 | NB69 | TACAGTCCGAGCCTCATGTGATCTATCTCGATCCCGCGAAATTAATACGACTCAC |
| F10 | NB70 | ACCGAGATCCTACGAATGGAGTGTATCTCGATCCCGCGAAATTAATACGACTCAC |
| F11 | NB71 | CCTGGGAGCATCAGGTAGTAACAGATCTCGATCCCGCGAAATTAATACGACTCAC |
| F12 | NB72 | TAGCTGACTGTCTTCCATACCGACATCTCGATCCCGCGAAATTAATACGACTCAC |
| G1 | NB73 | AAGAAACAGGATGACAGAACCCTCATCTCGATCCCGCGAAATTAATACGACTCAC |
| G2 | NB74 | TACAAGCATCCCAACACTTCCACTATCTCGATCCCGCGAAATTAATACGACTCAC |
| G3 | NB75 | GACCATTGTGATGAACCCTGTTGTATCTCGATCCCGCGAAATTAATACGACTCAC |

| | | |
|---|---|---|
| G4 | NB76 | ATGCTTGTTACATCAACCCTGGACATCTCGATCCCGCGAAATTAATACGACTCAC |
| G5 | NB77 | CGACCTGTTTCTCAGGGATACAACATCTCGATCCCGCGAAATTAATACGACTCAC |
| G6 | NB78 | AACAACCGAACCTTTGAATCAGAAATCTCGATCCCGCGAAATTAATACGACTCAC |
| G7 | NB79 | TCTCGGAGATAGTTCTCACTGCTGATCTCGATCCCGCGAAATTAATACGACTCAC |
| G8 | NB80 | CGGATGAACATAGGATAGCGATTCATCTCGATCCCGCGAAATTAATACGACTCAC |
| G9 | NB81 | CCTCATCTTGTGAAGTTGTTTCGGATCTCGATCCCGCGAAATTAATACGACTCAC |
| G10 | NB82 | ACGGTATGTCGAGTTCCAGGACTAATCTCGATCCCGCGAAATTAATACGACTCAC |
| G11 | NB83 | TGGCTTGATCTAGGTAAGGTCGAAATCTCGATCCCGCGAAATTAATACGACTCAC |
| G12 | NB84 | GTAGTGGACCTAGAACCTGTGCCAATCTCGATCCCGCGAAATTAATACGACTCAC |
| H1 | NB85 | AACGGAGGAGTTAGTTGGATGATCATCTCGATCCCGCGAAATTAATACGACTCAC |
| H2 | NB86 | AGGTGATCCCAACAAGCGTAAGTAATCTCGATCCCGCGAAATTAATACGACTCAC |
| H3 | NB87 | TACATGCTCCTGTTGTTAGGGAGGATCTCGATCCCGCGAAATTAATACGACTCAC |
| H4 | NB88 | TCTTCTACTACCGATCCGAAGCAGATCTCGATCCCGCGAAATTAATACGACTCAC |
| H5 | NB89 | ACAGCATCAATGTTTGGCTAGTTGATCTCGATCCCGCGAAATTAATACGACTCAC |
| H6 | NB90 | GATGTAGAGGGTACGGTTTGAGGCATCTCGATCCCGCGAAATTAATACGACTCAC |
| H7 | NB91 | GGCTCCATAGGAACTCACGCTACTATCTCGATCCCGCGAAATTAATACGACTCAC |
| H8 | NB92 | TTGTGAGTGGAAAGATACAGGACCATCTCGATCCCGCGAAATTAATACGACTCAC |
| H9 | NB93 | AGTTTCCATCACTTCAGACTTGGGATCTCGATCCCGCGAAATTAATACGACTCAC |
| H10 | NB94 | GATTGTCCTCAAACTGCCACCTACATCTCGATCCCGCGAAATTAATACGACTCAC |
| H11 | NB95 | CCTGTCTGGAAGAAGAATGGACTTATCTCGATCCCGCGAAATTAATACGACTCAC |
| H12 | NB96 | CTGAACGGTCATAGAGTCCACCATATCTCGATCCCGCGAAATTAATACGACTCAC |

**Table S4.** Full-length LevSeq reverse barcode-linked primer sequences used in this work. The primer-linked barcodes below are directly ordered from IDT.

| Well Position | Name | Full Length Order |
|---|---|---|
| A1 | RB01 | AAGAAAGTTGTCGGTGTCTTTGTGGCCCCAAGGGGTTATGCTAGTTATTGCTC |
| A2 | RB02 | TCGATTCCGTTTGTAGTCGTCTGTGCCCCAAGGGGTTATGCTAGTTATTGCTC |
| A3 | RB03 | GAGTCTTGTGTCCCAGTTACCAGGGCCCCAAGGGGTTATGCTAGTTATTGCTC |
| A4 | RB04 | TTCGGATTCTATCGTGTTTCCCTAGCCCCAAGGGGTTATGCTAGTTATTGCTC |
| A5 | RB05 | CTTGTCCAGGGTTTGTGTAACCTTGCCCCAAGGGGTTATGCTAGTTATTGCTC |
| A6 | RB06 | TTCTCGCAAAGGCAGAAAGTAGTCGCCCCAAGGGGTTATGCTAGTTATTGCTC |
| A7 | RB07 | GTGTTACCGTGGGAATGAATCCTTGCCCCAAGGGGTTATGCTAGTTATTGCTC |
| A8 | RB08 | TTCAGGGAACAAACCAAGTTACGTGCCCCAAGGGGTTATGCTAGTTATTGCTC |
| A9 | RB09 | AACTAGGCACAGCGAGTCTTGGTTGCCCCAAGGGGTTATGCTAGTTATTGCTC |
| A10 | RB10 | AAGCGTTGAAACCTTTGTCCTCTCGCCCCAAGGGGTTATGCTAGTTATTGCTC |
| A11 | RB11 | GTTTCATCTATCGGAGGGAATGGAGCCCCAAGGGGTTATGCTAGTTATTGCTC |
| A12 | RB12 | CAGGTAGAAAGAAGCAGAATCGGAGCCCCAAGGGGTTATGCTAGTTATTGCTC |
| B1 | RB13 | AGAACGACTTCCATACTCGTGTGAGCCCCAAGGGGTTATGCTAGTTATTGCTC |
| B2 | RB14 | AACGAGTCTCTTGGGACCCATAGAGCCCCAAGGGGTTATGCTAGTTATTGCTC |
| B3 | RB15 | AGGTCTACCTCGCTAACACCACTGGCCCCAAGGGGTTATGCTAGTTATTGCTC |
| B4 | RB16 | CGTCAACTGACAGTGGTTCGTACTGCCCCAAGGGGTTATGCTAGTTATTGCTC |
| B5 | RB17 | ACCCTCCAGGAAAGTACCTCTGATGCCCCAAGGGGTTATGCTAGTTATTGCTC |
| B6 | RB18 | CCAAACCCAACAACCTAGATAGGCGCCCCAAGGGGTTATGCTAGTTATTGCTC |
| B7 | RB19 | GTTCCTCGTGCAGTGTCAAGAGATGCCCCAAGGGGTTATGCTAGTTATTGCTC |
| B8 | RB20 | TTGCGTCCTGTTACGAGAACTCATGCCCCAAGGGGTTATGCTAGTTATTGCTC |
| B9 | RB21 | GAGCCTCTCATTGTCCGTTCTCTAGCCCCAAGGGGTTATGCTAGTTATTGCTC |
| B10 | RB22 | ACCACTGCCATGTATCAAAGTACGGCCCCAAGGGGTTATGCTAGTTATTGCTC |
| B11 | RB23 | CTTACTACCCAGTGAACCTCCTCGGCCCCAAGGGGTTATGCTAGTTATTGCTC |
| B12 | RB24 | GCATAGTTCTGCATGATGGGTTAGGCCCCAAGGGGTTATGCTAGTTATTGCTC |
| C1 | RB25 | GTAAGTTGGGTATGCAACGCAATGGCCCCAAGGGGTTATGCTAGTTATTGCTC |
| C2 | RB26 | CATACAGCGACTACGCATTCTCATGCCCCAAGGGGTTATGCTAGTTATTGCTC |
| C3 | RB27 | CGACGGTTAGATTCACCTCTTACAGCCCCAAGGGGTTATGCTAGTTATTGCTC |
| C4 | RB28 | TGAAACCTAAGAAGGCACCGTATCGCCCCAAGGGGTTATGCTAGTTATTGCTC |
| C5 | RB29 | CTAGACACCTTGGGTTGACAGACCGCCCCAAGGGGTTATGCTAGTTATTGCTC |
| C6 | RB30 | TCAGTGAGGATCTACTTCGACCCAGCCCCAAGGGGTTATGCTAGTTATTGCTC |
| C7 | RB31 | TGCGTACAGCAATCAGTTACATTGGCCCCAAGGGGTTATGCTAGTTATTGCTC |
| C8 | RB32 | CCAGTAGAAGTCCGACAACGTCATGCCCCAAGGGGTTATGCTAGTTATTGCTC |
| C9 | RB33 | CAGACTTGGTACGGTTGGGTAACTGCCCCAAGGGGTTATGCTAGTTATTGCTC |
| C10 | RB34 | GGACGAAGAACTCAAGTCAAAGGCGCCCCAAGGGGTTATGCTAGTTATTGCTC |
| C11 | RB35 | CTACTTACGAAGCTGAGGGACTGCGCCCCAAGGGGTTATGCTAGTTATTGCTC |
| C12 | RB36 | ATGTCCCAGTTAGAGGAGGAAACAGCCCCAAGGGGTTATGCTAGTTATTGCTC |

| | | |
|---|---|---|
| D1 | RB37 | GCTTGCGATTGATGCTTAGTATCAGCCCCAAGGGGTTATGCTAGTTATTGCTC |
| D2 | RB38 | ACCACAGGAGGACGATACAGAGAAGCCCCAAGGGGTTATGCTAGTTATTGCTC |
| D3 | RB39 | CCACAGTGTCAACTAGAGCCTCTCGCCCCAAGGGGTTATGCTAGTTATTGCTC |
| D4 | RB40 | TAGTTTGGATGACCAAGGATAGCCGCCCCAAGGGGTTATGCTAGTTATTGCTC |
| D5 | RB41 | GGAGTTCGTCCAGAGAAGTACACGGCCCCAAGGGGTTATGCTAGTTATTGCTC |
| D6 | RB42 | CTACGTGTAAGGCATACCTGCCAGGCCCCAAGGGGTTATGCTAGTTATTGCTC |
| D7 | RB43 | CTTTCGTTGTTGACTCGACGGTAGGCCCCAAGGGGTTATGCTAGTTATTGCTC |
| D8 | RB44 | AGTAGAAAGGGTTCCTTCCCACTCGCCCCAAGGGGTTATGCTAGTTATTGCTC |
| D9 | RB45 | GATCCAACAGAGATGCCTTCAGTGGCCCCAAGGGGTTATGCTAGTTATTGCTC |
| D10 | RB46 | GCTGTGTTCCACTTCATTCTCCTGGCCCCAAGGGGTTATGCTAGTTATTGCTC |
| D11 | RB47 | GTGCAACTTTCCCACAGGTAGTTCGCCCCAAGGGGTTATGCTAGTTATTGCTC |
| D12 | RB48 | CATCTGGAACGTGGTACACCTGTAGCCCCAAGGGGTTATGCTAGTTATTGCTC |
| E1 | RB49 | ACTGGTGCAGCTTTGAACATCTAGGCCCCAAGGGGTTATGCTAGTTATTGCTC |
| E2 | RB50 | ATGGACTTTGGTAACTTCCTGCGTGCCCCAAGGGGTTATGCTAGTTATTGCTC |
| E3 | RB51 | GTTGAATGAGCCTACTGGGTCCTCGCCCCAAGGGGTTATGCTAGTTATTGCTC |
| E4 | RB52 | TGAGAGACAAGATTGTTCGTGGACGCCCCAAGGGGTTATGCTAGTTATTGCTC |
| E5 | RB53 | AGATTCAGACCGTCTCATGCAAAGGCCCCAAGGGGTTATGCTAGTTATTGCTC |
| E6 | RB54 | CAAGAGCTTTGACTAAGGAGCATGGCCCCAAGGGGTTATGCTAGTTATTGCTC |
| E7 | RB55 | TGGAAGATGAGACCCTGATCTACGGCCCCAAGGGGTTATGCTAGTTATTGCTC |
| E8 | RB56 | TCACTACTCAACAGGTGGCATGAAGCCCCAAGGGGTTATGCTAGTTATTGCTC |
| E9 | RB57 | GCTAGGTCAATCTCCTTCGGAAGTGCCCCAAGGGGTTATGCTAGTTATTGCTC |
| E10 | RB58 | CAGGTTACTCCTCCGTGAGTCTGAGCCCCAAGGGGTTATGCTAGTTATTGCTC |
| E11 | RB59 | TCAATCAAGAAGGGAAAGCAAGGTGCCCCAAGGGGTTATGCTAGTTATTGCTC |
| E12 | RB60 | CATGTTCAACCAAGGCTTCTATGGGCCCCAAGGGGTTATGCTAGTTATTGCTC |
| F1 | RB61 | AGAGGGTACTATGTGCCTCAGCACGCCCCAAGGGGTTATGCTAGTTATTGCTC |
| F2 | RB62 | CACCCACACTTACTTCAGGACGTAGCCCCAAGGGGTTATGCTAGTTATTGCTC |
| F3 | RB63 | TTCTGAAGTTCCTGGGTCTTGAACGCCCCAAGGGGTTATGCTAGTTATTGCTC |
| F4 | RB64 | GACAGACACCGTTCATCGACTTTCGCCCCAAGGGGTTATGCTAGTTATTGCTC |
| F5 | RB65 | TTCTCAGTCTTCCTCCAGACAAGGGCCCCAAGGGGTTATGCTAGTTATTGCTC |
| F6 | RB66 | CCGATCCTTGTGGCTTCTAACTTCGCCCCAAGGGGTTATGCTAGTTATTGCTC |
| F7 | RB67 | GTTTGTCATACTCGTGTGCTCACCGCCCCAAGGGGTTATGCTAGTTATTGCTC |
| F8 | RB68 | GAATCTAAGCAAACACGAAGGTGGGCCCCAAGGGGTTATGCTAGTTATTGCTC |
| F9 | RB69 | TACAGTCCGAGCCTCATGTGATCTGCCCCAAGGGGTTATGCTAGTTATTGCTC |
| F10 | RB70 | ACCGAGATCCTACGAATGGAGTGTGCCCCAAGGGGTTATGCTAGTTATTGCTC |
| F11 | RB71 | CCTGGGAGCATCAGGTAGTAACAGGCCCCAAGGGGTTATGCTAGTTATTGCTC |
| F12 | RB72 | TAGCTGACTGTCTTCCATACCGACGCCCCAAGGGGTTATGCTAGTTATTGCTC |
| G1 | RB73 | AAGAAACAGGATGACAGAACCCTCGCCCCAAGGGGTTATGCTAGTTATTGCTC |
| G2 | RB74 | TACAAGCATCCCAACACTTCCACTGCCCCAAGGGGTTATGCTAGTTATTGCTC |
| G3 | RB75 | GACCATTGTGATGAACCCTGTTGTGCCCCAAGGGGTTATGCTAGTTATTGCTC |

| | | |
|---|---|---|
| G4 | RB76 | ATGCTTGTTACATCAACCCTGGACGCCCCAAGGGGTTATGCTAGTTATTGCTC |
| G5 | RB77 | CGACCTGTTTCTCAGGGATACAACGCCCCAAGGGGTTATGCTAGTTATTGCTC |
| G6 | RB78 | AACAACCGAACCTTTGAATCAGAAGCCCCAAGGGGTTATGCTAGTTATTGCTC |
| G7 | RB79 | TCTCGGAGATAGTTCTCACTGCTGGCCCCAAGGGGTTATGCTAGTTATTGCTC |
| G8 | RB80 | CGGATGAACATAGGATAGCGATTCGCCCCAAGGGGTTATGCTAGTTATTGCTC |
| G9 | RB81 | CCTCATCTTGTGAAGTTGTTTCGGGCCCCAAGGGGTTATGCTAGTTATTGCTC |
| G10 | RB82 | ACGGTATGTCGAGTTCCAGGACTAGCCCCAAGGGGTTATGCTAGTTATTGCTC |
| G11 | RB83 | TGGCTTGATCTAGGTAAGGTCGAAGCCCCAAGGGGTTATGCTAGTTATTGCTC |
| G12 | RB84 | GTAGTGGACCTAGAACCTGTGCCAGCCCCAAGGGGTTATGCTAGTTATTGCTC |
| H1 | RB85 | AACGGAGGAGTTAGTTGGATGATCGCCCCAAGGGGTTATGCTAGTTATTGCTC |
| H2 | RB86 | AGGTGATCCCAACAAGCGTAAGTAGCCCCAAGGGGTTATGCTAGTTATTGCTC |
| H3 | RB87 | TACATGCTCCTGTTGTTAGGGAGGGCCCCAAGGGGTTATGCTAGTTATTGCTC |
| H4 | RB88 | TCTTCTACTACCGATCCGAAGCAGGCCCCAAGGGGTTATGCTAGTTATTGCTC |
| H5 | RB89 | ACAGCATCAATGTTTGGCTAGTTGGCCCCAAGGGGTTATGCTAGTTATTGCTC |
| H6 | RB90 | GATGTAGAGGGTACGGTTTGAGGCGCCCCAAGGGGTTATGCTAGTTATTGCTC |
| H7 | RB91 | GGCTCCATAGGAACTCACGCTACTGCCCCAAGGGGTTATGCTAGTTATTGCTC |
| H8 | RB92 | TTGTGAGTGGAAAGATACAGGACCGCCCCAAGGGGTTATGCTAGTTATTGCTC |
| H9 | RB93 | AGTTTCCATCACTTCAGACTTGGGGCCCCAAGGGGTTATGCTAGTTATTGCTC |
| H10 | RB94 | GATTGTCCTCAAACTGCCACCTACGCCCCAAGGGGTTATGCTAGTTATTGCTC |
| H11 | RB95 | CCTGTCTGGAAGAAGAATGGACTTGCCCCAAGGGGTTATGCTAGTTATTGCTC |
| H12 | RB96 | CTGAACGGTCATAGAGTCCACCATGCCCCAAGGGGTTATGCTAGTTATTGCTC |

#### Plate Maps

This section contains all plate maps for the barcode plates (LevSeq plates) used in this study. The tables that follow show how the primers from the forward and reverse barcode plates (Tables S3 and S4) were arrayed to produce the barcode plates. Each entry in the below plate maps follows lists the forward primer on top and the reverse primer on the bottom. An "F" after the delimiter indicates that the barcode preceding the delimiter was from the forward barcode plate ("NB" in Table S3) and an "R" indicates that the well was from the reverse barcode plate ("RB" in Table S4). A detailed protocol for how the barcode plates were produced is given in *Preparation of LevSeq Barcode Primer Mixes*, above.

**Table S5.** Plate map for LevSeq01 used in this study.

| LevSeq01 | 01 | 02 | 03 | 04 | 05 | 06 | 07 | 08 | 09 | 10 | 11 | 12 |
|---|---|---|---|---|---|---|---|---|---|---|---|---|
| A | NB01-F<br>RB01-R | NB02-F<br>RB01-R | NB03-F<br>RB01-R | NB04-F<br>RB01-R | NB05-F<br>RB01-R | NB06-F<br>RB01-R | NB07-F<br>RB01-R | NB08-F<br>RB01-R | NB09-F<br>RB01-R | NB10-F<br>RB01-R | NB11-F<br>RB01-R | NB12-F<br>RB01-R |
| B | NB13-F<br>RB01-R | NB14-F<br>RB01-R | NB15-F<br>RB01-R | NB16-F<br>RB01-R | NB17-F<br>RB01-R | NB18-F<br>RB01-R | NB19-F<br>RB01-R | NB20-F<br>RB01-R | NB21-F<br>RB01-R | NB22-F<br>RB01-R | NB23-F<br>RB01-R | NB24-F<br>RB01-R |
| C | NB25-F<br>RB01-R | NB26-F<br>RB01-R | NB27-F<br>RB01-R | NB28-F<br>RB01-R | NB29-F<br>RB01-R | NB30-F<br>RB01-R | NB31-F<br>RB01-R | NB32-F<br>RB01-R | NB33-F<br>RB01-R | NB34-F<br>RB01-R | NB35-F<br>RB01-R | NB36-F<br>RB01-R |
| D | NB37-F<br>RB01-R | NB38-F<br>RB01-R | NB39-F<br>RB01-R | NB40-F<br>RB01-R | NB41-F<br>RB01-R | NB42-F<br>RB01-R | NB43-F<br>RB01-R | NB44-F<br>RB01-R | NB45-F<br>RB01-R | NB46-F<br>RB01-R | NB47-F<br>RB01-R | NB48-F<br>RB01-R |
| E | NB49-F<br>RB01-R | NB50-F<br>RB01-R | NB51-F<br>RB01-R | NB52-F<br>RB01-R | NB53-F<br>RB01-R | NB54-F<br>RB01-R | NB55-F<br>RB01-R | NB56-F<br>RB01-R | NB57-F<br>RB01-R | NB58-F<br>RB01-R | NB59-F<br>RB01-R | NB60-F<br>RB01-R |
| F | NB61-F<br>RB01-R | NB62-F<br>RB01-R | NB63-F<br>RB01-R | NB64-F<br>RB01-R | NB65-F<br>RB01-R | NB66-F<br>RB01-R | NB67-F<br>RB01-R | NB68-F<br>RB01-R | NB69-F<br>RB01-R | NB70-F<br>RB01-R | NB71-F<br>RB01-R | NB72-F<br>RB01-R |
| G | NB73-F<br>RB01-R | NB74-F<br>RB01-R | NB75-F<br>RB01-R | NB76-F<br>RB01-R | NB77-F<br>RB01-R | NB78-F<br>RB01-R | NB79-F<br>RB01-R | NB80-F<br>RB01-R | NB81-F<br>RB01-R | NB82-F<br>RB01-R | NB83-F<br>RB01-R | NB84-F<br>RB01-R |
| H | NB85-F<br>RB01-R | NB86-F<br>RB01-R | NB87-F<br>RB01-R | NB88-F<br>RB01-R | NB89-F<br>RB01-R | NB90-F<br>RB01-R | NB91-F<br>RB01-R | NB92-F<br>RB01-R | NB93-F<br>RB01-R | NB94-F<br>RB01-R | NB95-F<br>RB01-R | NB96-F<br>RB01-R |

**Table S6.** Plate map for LevSeq02 used in this study.

| LevSeq02 | 01 | 02 | 03 | 04 | 05 | 06 | 07 | 08 | 09 | 10 | 11 | 12 |
|---|---|---|---|---|---|---|---|---|---|---|---|---|
| A | NB01-F RB02-R | NB02-F RB02-R | NB03-F RB02-R | NB04-F RB02-R | NB05-F RB02-R | NB06-F RB02-R | NB07-F RB02-R | NB08-F RB02-R | NB09-F RB02-R | NB10-F RB02-R | NB11-F RB02-R | NB12-F RB02-R |
| B | NB13-F RB02-R | NB14-F RB02-R | NB15-F RB02-R | NB16-F RB02-R | NB17-F RB02-R | NB18-F RB02-R | NB19-F RB02-R | NB20-F RB02-R | NB21-F RB02-R | NB22-F RB02-R | NB23-F RB02-R | NB24-F RB02-R |
| C | NB25-F RB02-R | NB26-F RB02-R | NB27-F RB02-R | NB28-F RB02-R | NB29-F RB02-R | NB30-F RB02-R | NB31-F RB02-R | NB32-F RB02-R | NB33-F RB02-R | NB34-F RB02-R | NB35-F RB02-R | NB36-F RB02-R |
| D | NB37-F RB02-R | NB38-F RB02-R | NB39-F RB02-R | NB40-F RB02-R | NB41-F RB02-R | NB42-F RB02-R | NB43-F RB02-R | NB44-F RB02-R | NB45-F RB02-R | NB46-F RB02-R | NB47-F RB02-R | NB48-F RB02-R |
| E | NB49-F RB02-R | NB50-F RB02-R | NB51-F RB02-R | NB52-F RB02-R | NB53-F RB02-R | NB54-F RB02-R | NB55-F RB02-R | NB56-F RB02-R | NB57-F RB02-R | NB58-F RB02-R | NB59-F RB02-R | NB60-F RB02-R |
| F | NB61-F RB02-R | NB62-F RB02-R | NB63-F RB02-R | NB64-F RB02-R | NB65-F RB02-R | NB66-F RB02-R | NB67-F RB02-R | NB68-F RB02-R | NB69-F RB02-R | NB70-F RB02-R | NB71-F RB02-R | NB72-F RB02-R |
| G | NB73-F RB02-R | NB74-F RB02-R | NB75-F RB02-R | NB76-F RB02-R | NB77-F RB02-R | NB78-F RB02-R | NB79-F RB02-R | NB80-F RB02-R | NB81-F RB02-R | NB82-F RB02-R | NB83-F RB02-R | NB84-F RB02-R |
| H | NB85-F RB02-R | NB86-F RB02-R | NB87-F RB02-R | NB88-F RB02-R | NB89-F RB02-R | NB90-F RB02-R | NB91-F RB02-R | NB92-F RB02-R | NB93-F RB02-R | NB94-F RB02-R | NB95-F RB02-R | NB96-F RB02-R |

**Table S7.** Plate map for LevSeq03 used in this study.

| LevSeq03 | 01 | 02 | 03 | 04 | 05 | 06 | 07 | 08 | 09 | 10 | 11 | 12 |
|---|---|---|---|---|---|---|---|---|---|---|---|---|
| A | NB01-F RB03-R | NB02-F RB03-R | NB03-F RB03-R | NB04-F RB03-R | NB05-F RB03-R | NB06-F RB03-R | NB07-F RB03-R | NB08-F RB03-R | NB09-F RB03-R | NB10-F RB03-R | NB11-F RB03-R | NB12-F RB03-R |
| B | NB13-F RB03-R | NB14-F RB03-R | NB15-F RB03-R | NB16-F RB03-R | NB17-F RB03-R | NB18-F RB03-R | NB19-F RB03-R | NB20-F RB03-R | NB21-F RB03-R | NB22-F RB03-R | NB23-F RB03-R | NB24-F RB03-R |
| C | NB25-F RB03-R | NB26-F RB03-R | NB27-F RB03-R | NB28-F RB03-R | NB29-F RB03-R | NB30-F RB03-R | NB31-F RB03-R | NB32-F RB03-R | NB33-F RB03-R | NB34-F RB03-R | NB35-F RB03-R | NB36-F RB03-R |
| D | NB37-F RB03-R | NB38-F RB03-R | NB39-F RB03-R | NB40-F RB03-R | NB41-F RB03-R | NB42-F RB03-R | NB43-F RB03-R | NB44-F RB03-R | NB45-F RB03-R | NB46-F RB03-R | NB47-F RB03-R | NB48-F RB03-R |
| E | NB49-F RB03-R | NB50-F RB03-R | NB51-F RB03-R | NB52-F RB03-R | NB53-F RB03-R | NB54-F RB03-R | NB55-F RB03-R | NB56-F RB03-R | NB57-F RB03-R | NB58-F RB03-R | NB59-F RB03-R | NB60-F RB03-R |
| F | NB61-F RB03-R | NB62-F RB03-R | NB63-F RB03-R | NB64-F RB03-R | NB65-F RB03-R | NB66-F RB03-R | NB67-F RB03-R | NB68-F RB03-R | NB69-F RB03-R | NB70-F RB03-R | NB71-F RB03-R | NB72-F RB03-R |
| G | NB73-F RB03-R | NB74-F RB03-R | NB75-F RB03-R | NB76-F RB03-R | NB77-F RB03-R | NB78-F RB03-R | NB79-F RB03-R | NB80-F RB03-R | NB81-F RB03-R | NB82-F RB03-R | NB83-F RB03-R | NB84-F RB03-R |
| H | NB85-F RB03-R | NB86-F RB03-R | NB87-F RB03-R | NB88-F RB03-R | NB89-F RB03-R | NB90-F RB03-R | NB91-F RB03-R | NB92-F RB03-R | NB93-F RB03-R | NB94-F RB03-R | NB95-F RB03-R | NB96-F RB03-R |

**Table S8.** Plate map for LevSeq04 used in this study.

| LevSeq04 | 01 | 02 | 03 | 04 | 05 | 06 | 07 | 08 | 09 | 10 | 11 | 12 |
|---|---|---|---|---|---|---|---|---|---|---|---|---|
| A | NB01-F RB04-R | NB02-F RB04-R | NB03-F RB04-R | NB04-F RB04-R | NB05-F RB04-R | NB06-F RB04-R | NB07-F RB04-R | NB08-F RB04-R | NB09-F RB04-R | NB10-F RB04-R | NB11-F RB04-R | NB12-F RB04-R |
| B | NB13-F RB04-R | NB14-F RB04-R | NB15-F RB04-R | NB16-F RB04-R | NB17-F RB04-R | NB18-F RB04-R | NB19-F RB04-R | NB20-F RB04-R | NB21-F RB04-R | NB22-F RB04-R | NB23-F RB04-R | NB24-F RB04-R |
| C | NB25-F RB04-R | NB26-F RB04-R | NB27-F RB04-R | NB28-F RB04-R | NB29-F RB04-R | NB30-F RB04-R | NB31-F RB04-R | NB32-F RB04-R | NB33-F RB04-R | NB34-F RB04-R | NB35-F RB04-R | NB36-F RB04-R |
| D | NB37-F RB04-R | NB38-F RB04-R | NB39-F RB04-R | NB40-F RB04-R | NB41-F RB04-R | NB42-F RB04-R | NB43-F RB04-R | NB44-F RB04-R | NB45-F RB04-R | NB46-F RB04-R | NB47-F RB04-R | NB48-F RB04-R |
| E | NB49-F RB04-R | NB50-F RB04-R | NB51-F RB04-R | NB52-F RB04-R | NB53-F RB04-R | NB54-F RB04-R | NB55-F RB04-R | NB56-F RB04-R | NB57-F RB04-R | NB58-F RB04-R | NB59-F RB04-R | NB60-F RB04-R |
| F | NB61-F RB04-R | NB62-F RB04-R | NB63-F RB04-R | NB64-F RB04-R | NB65-F RB04-R | NB66-F RB04-R | NB67-F RB04-R | NB68-F RB04-R | NB69-F RB04-R | NB70-F RB04-R | NB71-F RB04-R | NB72-F RB04-R |
| G | NB73-F RB04-R | NB74-F RB04-R | NB75-F RB04-R | NB76-F RB04-R | NB77-F RB04-R | NB78-F RB04-R | NB79-F RB04-R | NB80-F RB04-R | NB81-F RB04-R | NB82-F RB04-R | NB83-F RB04-R | NB84-F RB04-R |
| H | NB85-F RB04-R | NB86-F RB04-R | NB87-F RB04-R | NB88-F RB04-R | NB89-F RB04-R | NB90-F RB04-R | NB91-F RB04-R | NB92-F RB04-R | NB93-F RB04-R | NB94-F RB04-R | NB95-F RB04-R | NB96-F RB04-R |

**Table S9.** Plate map for LevSeq05 used in this study.

| LevSeq05 | 01 | 02 | 03 | 04 | 05 | 06 | 07 | 08 | 09 | 10 | 11 | 12 |
|---|---|---|---|---|---|---|---|---|---|---|---|---|
| A | NB01-F RB05-R | NB02-F RB05-R | NB03-F RB05-R | NB04-F RB05-R | NB05-F RB05-R | NB06-F RB05-R | NB07-F RB05-R | NB08-F RB05-R | NB09-F RB05-R | NB10-F RB05-R | NB11-F RB05-R | NB12-F RB05-R |
| B | NB13-F RB05-R | NB14-F RB05-R | NB15-F RB05-R | NB16-F RB05-R | NB17-F RB05-R | NB18-F RB05-R | NB19-F RB05-R | NB20-F RB05-R | NB21-F RB05-R | NB22-F RB05-R | NB23-F RB05-R | NB24-F RB05-R |
| C | NB25-F RB05-R | NB26-F RB05-R | NB27-F RB05-R | NB28-F RB05-R | NB29-F RB05-R | NB30-F RB05-R | NB31-F RB05-R | NB32-F RB05-R | NB33-F RB05-R | NB34-F RB05-R | NB35-F RB05-R | NB36-F RB05-R |
| D | NB37-F RB05-R | NB38-F RB05-R | NB39-F RB05-R | NB40-F RB05-R | NB41-F RB05-R | NB42-F RB05-R | NB43-F RB05-R | NB44-F RB05-R | NB45-F RB05-R | NB46-F RB05-R | NB47-F RB05-R | NB48-F RB05-R |
| E | NB49-F RB05-R | NB50-F RB05-R | NB51-F RB05-R | NB52-F RB05-R | NB53-F RB05-R | NB54-F RB05-R | NB55-F RB05-R | NB56-F RB05-R | NB57-F RB05-R | NB58-F RB05-R | NB59-F RB05-R | NB60-F RB05-R |
| F | NB61-F RB05-R | NB62-F RB05-R | NB63-F RB05-R | NB64-F RB05-R | NB65-F RB05-R | NB66-F RB05-R | NB67-F RB05-R | NB68-F RB05-R | NB69-F RB05-R | NB70-F RB05-R | NB71-F RB05-R | NB72-F RB05-R |
| G | NB73-F RB05-R | NB74-F RB05-R | NB75-F RB05-R | NB76-F RB05-R | NB77-F RB05-R | NB78-F RB05-R | NB79-F RB05-R | NB80-F RB05-R | NB81-F RB05-R | NB82-F RB05-R | NB83-F RB05-R | NB84-F RB05-R |
| H | NB85-F RB05-R | NB86-F RB05-R | NB87-F RB05-R | NB88-F RB05-R | NB89-F RB05-R | NB90-F RB05-R | NB91-F RB05-R | NB92-F RB05-R | NB93-F RB05-R | NB94-F RB05-R | NB95-F RB05-R | NB96-F RB05-R |

**Table S10.** Plate map for LevSeq06 used in this study.

| LevSeq06 | 01 | 02 | 03 | 04 | 05 | 06 | 07 | 08 | 09 | 10 | 11 | 12 |
|---|---|---|---|---|---|---|---|---|---|---|---|---|
| A | NB01-F RB06-R | NB02-F RB06-R | NB03-F RB06-R | NB04-F RB06-R | NB05-F RB06-R | NB06-F RB06-R | NB07-F RB06-R | NB08-F RB06-R | NB09-F RB06-R | NB10-F RB06-R | NB11-F RB06-R | NB12-F RB06-R |
| B | NB13-F RB06-R | NB14-F RB06-R | NB15-F RB06-R | NB16-F RB06-R | NB17-F RB06-R | NB18-F RB06-R | NB19-F RB06-R | NB20-F RB06-R | NB21-F RB06-R | NB22-F RB06-R | NB23-F RB06-R | NB24-F RB06-R |
| C | NB25-F RB06-R | NB26-F RB06-R | NB27-F RB06-R | NB28-F RB06-R | NB29-F RB06-R | NB30-F RB06-R | NB31-F RB06-R | NB32-F RB06-R | NB33-F RB06-R | NB34-F RB06-R | NB35-F RB06-R | NB36-F RB06-R |
| D | NB37-F RB06-R | NB38-F RB06-R | NB39-F RB06-R | NB40-F RB06-R | NB41-F RB06-R | NB42-F RB06-R | NB43-F RB06-R | NB44-F RB06-R | NB45-F RB06-R | NB46-F RB06-R | NB47-F RB06-R | NB48-F RB06-R |
| E | NB49-F RB06-R | NB50-F RB06-R | NB51-F RB06-R | NB52-F RB06-R | NB53-F RB06-R | NB54-F RB06-R | NB55-F RB06-R | NB56-F RB06-R | NB57-F RB06-R | NB58-F RB06-R | NB59-F RB06-R | NB60-F RB06-R |
| F | NB61-F RB06-R | NB62-F RB06-R | NB63-F RB06-R | NB64-F RB06-R | NB65-F RB06-R | NB66-F RB06-R | NB67-F RB06-R | NB68-F RB06-R | NB69-F RB06-R | NB70-F RB06-R | NB71-F RB06-R | NB72-F RB06-R |
| G | NB73-F RB06-R | NB74-F RB06-R | NB75-F RB06-R | NB76-F RB06-R | NB77-F RB06-R | NB78-F RB06-R | NB79-F RB06-R | NB80-F RB06-R | NB81-F RB06-R | NB82-F RB06-R | NB83-F RB06-R | NB84-F RB06-R |
| H | NB85-F RB06-R | NB86-F RB06-R | NB87-F RB06-R | NB88-F RB06-R | NB89-F RB06-R | NB90-F RB06-R | NB91-F RB06-R | NB92-F RB06-R | NB93-F RB06-R | NB94-F RB06-R | NB95-F RB06-R | NB96-F RB06-R |

**Table S11.** Plate map for LevSeq07 used in this study.

| LevSeq07 | 01 | 02 | 03 | 04 | 05 | 06 | 07 | 08 | 09 | 10 | 11 | 12 |
|---|---|---|---|---|---|---|---|---|---|---|---|---|
| A | NB01-F RB07-R | NB02-F RB07-R | NB03-F RB07-R | NB04-F RB07-R | NB05-F RB07-R | NB06-F RB07-R | NB07-F RB07-R | NB08-F RB07-R | NB09-F RB07-R | NB10-F RB07-R | NB11-F RB07-R | NB12-F RB07-R |
| B | NB13-F RB07-R | NB14-F RB07-R | NB15-F RB07-R | NB16-F RB07-R | NB17-F RB07-R | NB18-F RB07-R | NB19-F RB07-R | NB20-F RB07-R | NB21-F RB07-R | NB22-F RB07-R | NB23-F RB07-R | NB24-F RB07-R |
| C | NB25-F RB07-R | NB26-F RB07-R | NB27-F RB07-R | NB28-F RB07-R | NB29-F RB07-R | NB30-F RB07-R | NB31-F RB07-R | NB32-F RB07-R | NB33-F RB07-R | NB34-F RB07-R | NB35-F RB07-R | NB36-F RB07-R |
| D | NB37-F RB07-R | NB38-F RB07-R | NB39-F RB07-R | NB40-F RB07-R | NB41-F RB07-R | NB42-F RB07-R | NB43-F RB07-R | NB44-F RB07-R | NB45-F RB07-R | NB46-F RB07-R | NB47-F RB07-R | NB48-F RB07-R |
| E | NB49-F RB07-R | NB50-F RB07-R | NB51-F RB07-R | NB52-F RB07-R | NB53-F RB07-R | NB54-F RB07-R | NB55-F RB07-R | NB56-F RB07-R | NB57-F RB07-R | NB58-F RB07-R | NB59-F RB07-R | NB60-F RB07-R |
| F | NB61-F RB07-R | NB62-F RB07-R | NB63-F RB07-R | NB64-F RB07-R | NB65-F RB07-R | NB66-F RB07-R | NB67-F RB07-R | NB68-F RB07-R | NB69-F RB07-R | NB70-F RB07-R | NB71-F RB07-R | NB72-F RB07-R |
| G | NB73-F RB07-R | NB74-F RB07-R | NB75-F RB07-R | NB76-F RB07-R | NB77-F RB07-R | NB78-F RB07-R | NB79-F RB07-R | NB80-F RB07-R | NB81-F RB07-R | NB82-F RB07-R | NB83-F RB07-R | NB84-F RB07-R |
| H | NB85-F RB07-R | NB86-F RB07-R | NB87-F RB07-R | NB88-F RB07-R | NB89-F RB07-R | NB90-F RB07-R | NB91-F RB07-R | NB92-F RB07-R | NB93-F RB07-R | NB94-F RB07-R | NB95-F RB07-R | NB96-F RB07-R |

**Table S12.** Plate map for LevSeq08 used in this study.

| LevSeq08 | 01 | 02 | 03 | 04 | 05 | 06 | 07 | 08 | 09 | 10 | 11 | 12 |
|---|---|---|---|---|---|---|---|---|---|---|---|---|
| A | NB01-F RB08-R | NB02-F RB08-R | NB03-F RB08-R | NB04-F RB08-R | NB05-F RB08-R | NB06-F RB08-R | NB07-F RB08-R | NB08-F RB08-R | NB09-F RB08-R | NB10-F RB08-R | NB11-F RB08-R | NB12-F RB08-R |
| B | NB13-F RB08-R | NB14-F RB08-R | NB15-F RB08-R | NB16-F RB08-R | NB17-F RB08-R | NB18-F RB08-R | NB19-F RB08-R | NB20-F RB08-R | NB21-F RB08-R | NB22-F RB08-R | NB23-F RB08-R | NB24-F RB08-R |
| C | NB25-F RB08-R | NB26-F RB08-R | NB27-F RB08-R | NB28-F RB08-R | NB29-F RB08-R | NB30-F RB08-R | NB31-F RB08-R | NB32-F RB08-R | NB33-F RB08-R | NB34-F RB08-R | NB35-F RB08-R | NB36-F RB08-R |
| D | NB37-F RB08-R | NB38-F RB08-R | NB39-F RB08-R | NB40-F RB08-R | NB41-F RB08-R | NB42-F RB08-R | NB43-F RB08-R | NB44-F RB08-R | NB45-F RB08-R | NB46-F RB08-R | NB47-F RB08-R | NB48-F RB08-R |
| E | NB49-F RB08-R | NB50-F RB08-R | NB51-F RB08-R | NB52-F RB08-R | NB53-F RB08-R | NB54-F RB08-R | NB55-F RB08-R | NB56-F RB08-R | NB57-F RB08-R | NB58-F RB08-R | NB59-F RB08-R | NB60-F RB08-R |
| F | NB61-F RB08-R | NB62-F RB08-R | NB63-F RB08-R | NB64-F RB08-R | NB65-F RB08-R | NB66-F RB08-R | NB67-F RB08-R | NB68-F RB08-R | NB69-F RB08-R | NB70-F RB08-R | NB71-F RB08-R | NB72-F RB08-R |
| G | NB73-F RB08-R | NB74-F RB08-R | NB75-F RB08-R | NB76-F RB08-R | NB77-F RB08-R | NB78-F RB08-R | NB79-F RB08-R | NB80-F RB08-R | NB81-F RB08-R | NB82-F RB08-R | NB83-F RB08-R | NB84-F RB08-R |
| H | NB85-F RB08-R | NB86-F RB08-R | NB87-F RB08-R | NB88-F RB08-R | NB89-F RB08-R | NB90-F RB08-R | NB91-F RB08-R | NB92-F RB08-R | NB93-F RB08-R | NB94-F RB08-R | NB95-F RB08-R | NB96-F RB08-R |

**Table S13.** Primers specific to the backbone of pET22b(+) used in *Par*LQ error prone library construction.

| Name | Direction | Sequence |
|---|---|---|
| 005 | Forward | GAAATAATTTTGTTTAACTTTAAGAAGGAGATATACATATG |
| 006 | Reverse | CAGTGCTAGGTGAAGGAATACCGCCAAGCGGAA |

**Table S14.** The LevSeq barcode plates used for sequencing *Par*LQ error prone mutagenesis libraries.

| Library ID | Barcode plate |
|---|---|
| *Par*LQ-ep1 | LevSeq01 |
| *Par*LQ-ep2 | LevSeq02 |
| *Par*LQ-ep3 | LevSeq03 |
| *Par*LQ-ep4 | LevSeq04 |
| *Par*LQ-ep5 | LevSeq01 |
| *Par*LQ-ep6 | LevSeq02 |
| *Par*LQ-ep7 | LevSeq03 |
| *Par*LQ-ep8 | LevSeq05 |
| *Par*LQ-ep9 | LevSeq06 |
| *Par*LQ-ep10 | LevSeq07 |
